# Dissecting reprogramming heterogeneity at single-cell resolution using scTF-seq

**DOI:** 10.1101/2024.01.30.577921

**Authors:** Wangjie Liu, Wouter Saelens, Pernille Rainer, Marjan Biočanin, Vincent Gardeux, Antoni Gralak, Guido van Mierlo, Julie Russeil, Tingdang Liu, Wanze Chen, Bart Deplancke

**Author notes:** Correspondence (W.C), (B.D). First author (Wangjie Liu, Wouter Saelens, Pernille Rainer).

## Abstract

Reprogramming approaches often produce heterogeneous cell fates and the mechanisms behind this heterogeneity are not well-understood. To address this gap, we developed scTF-seq, a technique inducing single-cell barcoded and doxycycline-inducible TF overexpression while quantifying TF dose-dependent transcriptomic changes. Applied to mouse embryonic multipotent stromal cells (MSCs), scTF-seq produced a gain-of-function atlas for 384 murine TFs. This atlas offers a valuable resource for gene regulation and reprogramming research, identifying key TFs governing MSC lineage differentiation, cell cycle control, and their interplay. Leveraging the single-cell resolution, we dissected reprogramming heterogeneity along dose and pseudotime. We thereby revealed TF dose-dependent and stochastic cell fate branching, unveiling gene expression signatures that enhance our understanding and prediction of reprogramming efficiency. scTF-seq also allowed us to classify TFs into four sensitivity classes based on dose response and determining features. Finally, in combinatorial scTF-seq, we observed that the same TF can exhibit both synergistic and antagonistic effects on another TF depending on its dose. In summary, scTF-seq provides a powerful tool for gaining mechanistic insights into how TFs determine cell states, while offering novel perspectives for cellular engineering strategies.

## Introduction

Understanding and controlling cell fates through gene regulatory programs, and particularly via transcription factor (TF)-mediated cell reprogramming, are critical objectives in biomedical research. Past studies employing the ectopic expression of single TFs or combinations have identified ‘master regulators’ that influence various cellular processes^1–3^, including differentiation, transdifferentiation, dedifferentiation, and reprogramming^4^. Here, we collectively refer to these processes as cell “reprogramming”. For instance, the “Yamanaka factors” (OCT3/4, SOX2, KLF4, and c-MYC) can convert adult fibroblasts into induced pluripotent stem cells (iPSCs)^5,6^.

However, reprogramming is typically characterized by significant heterogeneity and inefficiency, posing a major challenge^4,7–9^. This reprogramming heterogeneity is not solely due to cell-to-cell variability of the starting population^9,10^, as for example advancements in single-cell technology have revealed that cells can follow multiple branches along the path of reprogramming^11^. In addition, inhibition of proliferation or cell cycle synchronization significantly increased the efficiency of reprogramming, highlighting an important role for the cell cycle in modulating the capacity of a cell to be reprogrammed^12^. Nevertheless, it remains poorly understood which molecular mechanisms underlie cell fate branching and TF-cell cycle interaction during cell reprogramming. Another aspect that has historically received relatively little attention is the role of TF dose, even though TFs are known to vary in copy number over several orders of magnitude^13^. The dose of a TF does not only affect gene expression levels but also the set of targeted genes^13–15^. Consequently, TF dose may equally be key in steering cell reprogramming and thus account for the observed heterogeneity. The multifaceted nature of reprogramming is one of the principal reasons why it remains challenging to collectively study heterogeneity-contributing factors and how they influence cell reprogramming, especially when using bulk assays which are constrained by population-averaging readouts.

To answer these questions, a systematic quantitative TF screen at the single-cell level is essential to link TF function to reprogramming efficiency. TF overexpression would thereby be preferred since it can induce cell differentiation more efficiently than CRISPR activation^16,17^. In the past five years, several studies have implemented TF overexpression screens by coupling pooled TF overexpression with high-throughput readouts of single-cell RNA sequencing (scRNA-seq) or single-cell multiomics^16,18–20^. However, none have systematically investigated the roles of TF dose, cell cycle, and their interplay in steering cell reprogramming. To address this gap, we developed scTF-seq, aligning doxycycline-inducible barcoded overexpression of individual TFs with transcriptomic changes captured by scRNA-seq. This allowed us to map reprogramming properties of each TF and its dose at single-cell resolution. We conducted scTF-seq on mouse embryonic multipotent stromal cells (MSCs) for 419 mouse TFs in parallel. After rigid quality controls, this yielded a high-quality dataset that tabulates the TF overexpression level and respective TF-induced transcriptomic change for each of 39,187 cells linked to 384 TFs and 7 TF pairs. Our approaches identified novel cell reprogramming capacities of both known and uncharacterized TFs. Additionally, we were able to systematically study heterogeneous molecular and cellular responses resulting from TF dose, stochasticity, and/or cell cycle dynamics. Finally, targeted combinatorial TF analysis revealed that the same combination of TFs can interact synergistically and antagonistically depending on the TF dose. Our TF overexpression clone library, single-cell TF gain-of-function atlas, and analytic frameworks serve as valuable resources for achieving a mechanistic understanding of the roles of TFs in governing cell states.

## Results

### Constructing the scTF-seq library and single-cell atlas

To establish scTF-seq, we first built a lentiviral expression library for the open reading frames (ORFs) of 419 TFs (**Fig. 1a**, **Supplementary Table 1**, and **Methods**). Each ORF was associated with a unique barcode (termed TF-ID hereafter) close to the 3’ UTR, enabling precise identification of TFs and their respective expression levels by 3’ transcript sequencing, while preventing ORF length-dependent bias during PCR amplification. To control both the timing and dose of TF expression, we integrated a doxycycline-inducible promoter in the overexpression construct (**Fig. 1a, b**). Importantly, all viral particles were produced by individually packaging each vector to avoid barcode recombination and ensure more efficient and controllable TF expression than pooled virus packaging as used in most published screens^3,16,18–20^. In downstream virus transduction, our library can be used for both pooled and arrayed TF overexpression screening, serving as a resource for studying TFs of interest.

**Fig. 1:**
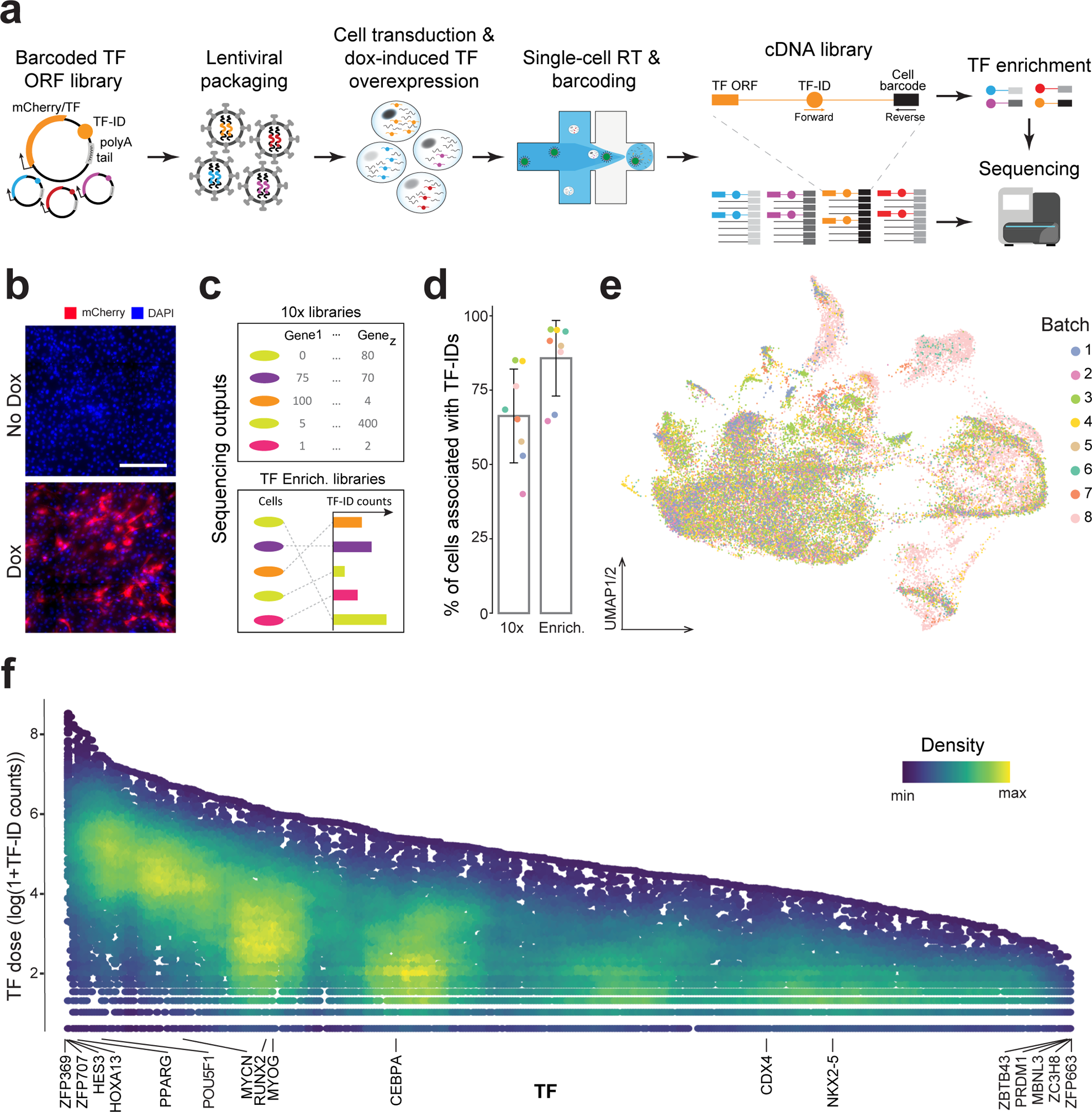
Overviews of scTF-seq design and TF overexpression atlas. **a** Schematic of the scTF-seq workflow. TF-ID, a unique barcode designed for mCherry (as control) or each individual TF; ORF, open reading frame; dox, doxycycline; RT, reverse transcription; Forward and Reverse, primers to enrich TF-IDs. **b** Representative fluorescent images of mCherry (red) and nuclei (DAPI, blue) on C3H10T1/2 cells treated without (No Dox) or with doxycycline (Dox). Scale bar, 125 μm. c Schematic of the sequencing outputs of scTF-seq: count matrices of gene expression in 10x libraries (top) and ectopic TF-ID expression in TF enrichment libraries (bottom) for each sequenced cell. **d** Percentage of cell barcodes associated with TF-IDs in 10x or TF enrichment libraries. Colors represent experiments (also referred to as “batches”, see color legend in **e**). **e** Uniform manifold approximation and projection (UMAP) of scRNA-seq data from 39,187 cells overexpressing 384 TFs after quality control and preprocessing (referred to as “TF atlas”). Colors represent batches. **f** Log-transformed TF expression levels (TF dose) of cells overexpressing individual TFs. Colors indicate cell density (number of neighbors) after randomly sampling up to 500 cells for each TF. See also **Supplementary Fig. 1**.

To assess the functionality of the scTF-seq library, we introduced it into mouse MSCs (C3H10T1/2)^21^ by lentiviral transduction in an arrayed manner (**Fig. 1a**) to achieve high transduction efficiencies and overexpression of individual TFs. We chose C3H10T1/2 cells due to their multipotent stem cell nature, enabling them to differentiate into adipocytes, chondrocytes, osteoblasts, or myocytes, thus providing a diverse range of cell fates to investigate the reprogramming capacity of TFs^22–24^. The expression of barcoded TFs was then induced by doxycycline treatment in a basic culture medium. To correct for the potential change of state and spontaneous differentiation of C3H10T1/2 cells when reaching confluence^21,25^, C3H10T1/2 cells that overexpressed barcoded mCherry under non-confluent and confluent conditions were employed as controls. In addition, we specifically treated control cells with an adipogenic cocktail (**Methods**) or induced *Myog* (encoding a key myogenesis regulator^26^) overexpression in them for up to 5 days to establish reference populations of mature adipocytes (called ‘Adipo_ref’) and myocytes (called ‘Myo_ref’). The transcriptomes of all cells were profiled using a droplet-based scRNA-seq approach (**Methods**). To ensure a high TF-ID recovery rate, we implemented an additional amplification and sequencing step targeting TF-IDs and cell barcodes (**Fig. 1a, c**, **Methods**), which increased the percentage of cell barcodes associated with a TF-ID from 67% to 86% (**Fig. 1d**, and **Supplementary Table 2**). Cells without any TF-ID were filtered out. The number of reads aligning to the overexpression construct exhibited a robust correlation between the TF enrichment and the conventional single-cell libraries (**Supplementary Fig. 1a**). We thus used TF-enrichment libraries to assign TFs to cells. To limit the risk of wrongly assigning TFs due to sequencing errors, only TF-IDs with a hamming distance of at least 2 nts were used within the same experiment (**Supplementary Fig. 1b**). Cell doublets were removed based on the proportion of different TF-IDs detected in each cell (**Supplementary Fig. 1c** and **Methods**). After further filtering low quality cells and TFs with too few assigned cells (**Supplementary Fig. 1d**), we obtained 39,187 cells covering 384 individual TFs and 7 TF pairs (detailed in the following sections). The number of cells (on average 100 cells per TF or TF pair) was uniformly distributed among TFs and experiments, supporting the advantage of array-based sample preparation (**Fig. 1e** and **Supplementary Fig. 1e**). By utilizing the TF enrichment library as an accurate readout of the TF-ID, we quantified the relative TF overexpression level in a cell by the log-transformed UMI count of its assigned TF-ID (referred to from now on as “TF dose”). In line with the design of scTF-seq, we observed variation in TF dose across cells for most TFs (**Fig. 1f**). Thus, TF dose functions here as an mRNA-based proxy for TF protein levels, a sensible approach given the generally reasonable correlation between protein and mRNA abundance across various contexts^27^.

### Identifying TFs directing lineage differentiation

Given that the activation of lineage developmental genes generally occurs in the G0/G1 phase^28^, we first focused on G0/G1 cells (**Supplementary Fig. 1f**, **Fig. 5a** and **Methods**) to study the roles of TFs in directing lineage differentiation. For each TF-control pair, we measured the main TF-driven transcriptomic variation by calculating the Euclidean distance of cells reprogrammed by TFs (simplified as “TF cells” hereafter) to the centroids of control cells in the space consisting of significant principal component coordinates (**Supplementary Fig. 2a-c** and **Methods**). We observed that the distances between certain TF cells and the centroids of control cells are negligible (**Supplementary Fig. 2b**). We labeled these cells as ‘non-functional’ ones given the close resemblance between the transcriptomes of those TF cells and control cells. This phenomenon was observed for many TFs, but typically only in a subset of TF cells, implying that TF overexpression tends to induce various degrees of transcriptomic alterations. Upon closer inspection, we found that higher doses correlate with more pronounced transcriptional changes, indicating that TF dose is a primary determinant of this reprogramming heterogeneity (**Supplementary Fig. 2c**). To mitigate dose variation and the potential interference arising from the ‘non-functional’ cells, we established a common threshold for all TFs. Through different iterations, we determined that the retention of ‘functional’ cells and exclusion of ‘non-functional’ ones is balanced at the 80th percentile of the distance between control cells to their centroids (**Supplementary Fig. 2b, d, and e**). Subsequently, we performed a clustering analysis of the atlas without non-functional TF cells, while retaining control cells (**Fig. 2a, b** and **Supplementary Fig. 2f**). Clusters 2, 4, and 6 expressed strikingly higher levels of differentiation genes such as *Bglap2*, *Fabp4,* and *Mylpf* (**Supplementary Fig. 2g**), representing osteogenic, adipogenic, and myogenic lineages, respectively. Adipogenic and myogenic reference cells (see above) co-localized with cluster 4 and 6 (**Fig. 2a, b**), validating the respectively adipogenic or myogenic nature of the two clusters. Cluster 7 exhibited high expression of interferon-stimulated genes like *Isg15* (**Supplementary Fig. 2g**), as well as a more general enrichment for inflammatory response as revealed by gene set enrichment analysis (GSEA) (**Supplementary Fig. 2h**). Cells reprogrammed by HEY1^29^, LZTS2^30^, HNF4A^31^, ZFP24, and ZFP692 were predominantly distributed in cluster 7. Despite the lack of clear functional information associated with inflammation for these TFs, the co-localization of their cells in cluster 7 with IRF3 (encoding a well-established immunomodulator^32^) cells suggests an involvement of these TFs in regulating inflammatory response genes.

**Fig. 2:**
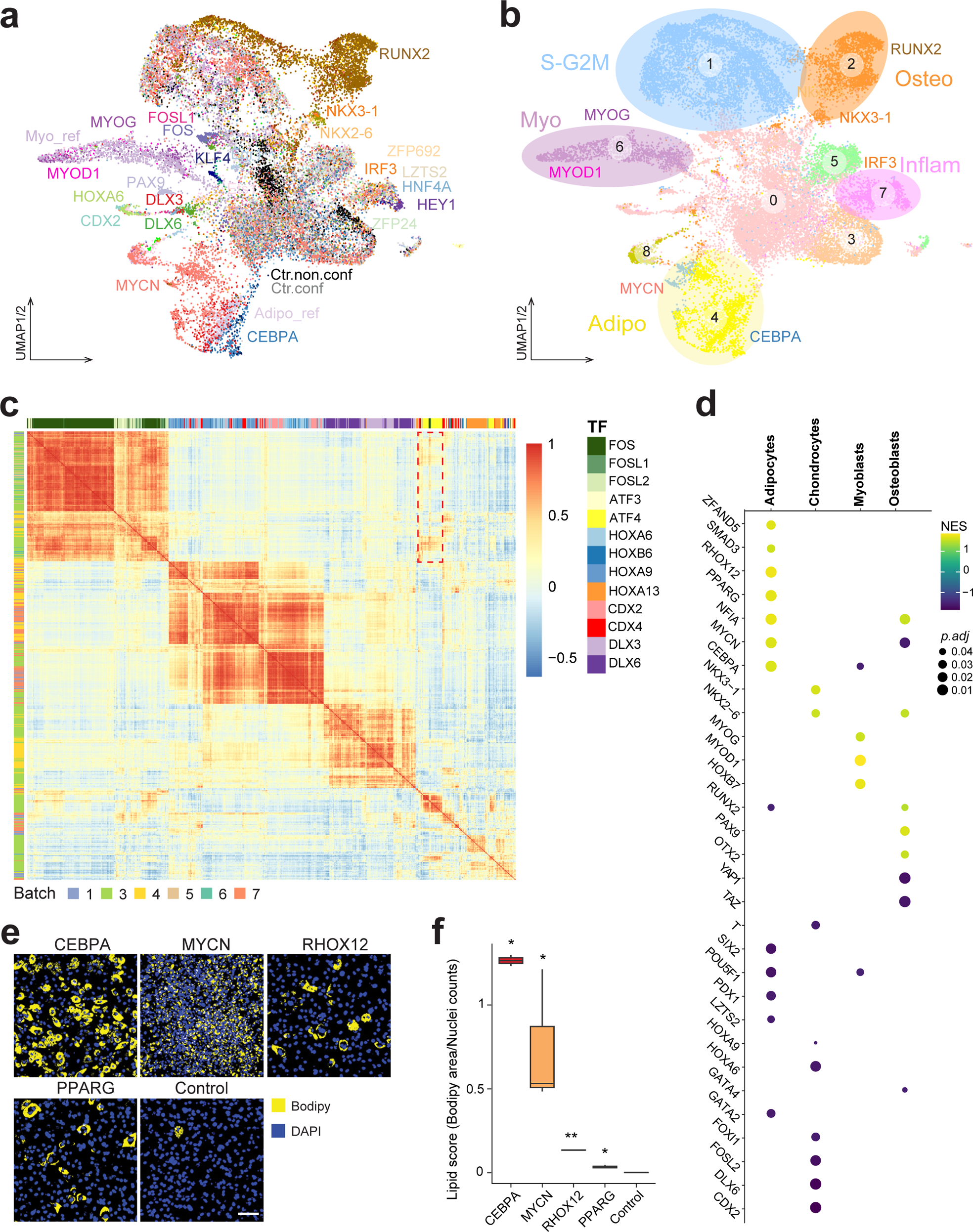
TFs directing lineage differentiation and immunomodulation. **a, b** UMAP of the integrated TF atlas with control, functional, and proliferating cells (referred to as the “functional TF atlas”). Colors indicate assigned TFs (**a**) and clusters (**b**). Ctr.conf and Ctr.non.conf in **a** represent confluent and non-confluent control (mCherry-overexpressing) cells, respectively. Colored circles in **b** highlight clusters having gene expression profiles related to myogenic, osteogenic or adipogenic lineages, S-G2M phase or immunomodulation. **c** Heatmap showing a pairwise Pearson correlation of functional TF cells annotated by TF (in column) and batch (in row). Cells are ordered by hierarchical clustering. **d** Dot plot showing a functional cell expression profile enrichment of each TF in the four main differentiation lineages of multipotent stromal stem cells. Only TFs having at least 25 functional cells and enriched in at least one of the four lineages with adjusted *P* value < 0.05 are shown. **e** Representative fluorescent images of lipids droplets (stained with Bodipy, yellow) and nuclei (stained with DAPI, blue) in C3H10T1/2 cells after 5 days of doxycycline-induced *Cebpa*, *Mycn*, *Rhox12*, *Pparg* and mCherry (control) overexpression. Scale bar, 100 µm. **f** Box plot showing lipid scores (quantified by Bodipy area/Nuclei counts on the images shown in panel **e**) of individual TFs and the control. At least 3 biological replicates for each. * *P* value < 0.05, ** *P* value < 0.01, pairwise two-sided *t*-test followed by FDR correction. See also **Supplementary Fig. 2**.

Next, we inferred functional TF modules governing the same gene expression programs by computing the transcriptomic similarity between TF cells (**Methods**). As depicted in **Fig. 2c** and **Supplementary Fig. 2i**, we found that homologous TFs can display varying degrees of similarity, while TFs from different families may still possess analogous functions. For instance, pronounced correlations were detected in both intra- and inter-families among *Cdx*, *Hox*, and *Dlx* families which regulate anterior-posterior patterning (**Fig. 2c**). Evolutionarily, the *Cdx*, *Hox*, and *Dlx* genes are also interrelated, all tracing back to a shared ancestral gene^33^. However, the correlation was particularly strong among HOXA6, HOXA9, HOXB6, CDX2, and CDX4 cells, but was less evident with cells reprogrammed by HOXA13 (**Fig. 2c**), corroborating the distinct role for HOXA13, as delineated in prior research^34,35^. In addition, we found that analogous functional characteristics are also consistent with known physical interactions among TFs such as the case for cells reprogrammed by FOS, FOSL1, and FOSL2, ATF3, and ATF4 (**Fig. 2c**). These *Fos* and *Atf* family-related TFs form cross-family heterodimers commonly referred to as activator protein 1 (AP-1)^36^, rationalizing why they induce similar transcriptomic responses. Thus, these results suggest that our scTF-seq atlas constitutes a valuable resource to explore the interaction between TFs, such as direct regulation and functional similarity.

In order to identify TFs that drive the differentiation of the four central MSC lineages, we performed GSEA analyses. These revealed various TFs whose differentially expressed genes are either positively or negatively enriched for expressed gene sets of osteoblasts, myoblasts, chondrocytes, and adipocytes (**Fig. 2d**). Many TFs induced a lineage expression program consistent with their previously reported role such as RUNX2 and PAX9 for osteogenesis^37,38^; HOXB7, MYOG, and MYOD1 for myogenesis^39,40^; NKX3-1 for chondrogenesis^41^; and SMAD3, PPARG, and CEBPA for adipogenesis^42–44^ (**Fig. 2d**). We also identified novel TF candidates that might be implicated in MSC lineages (**Fig. 2d**), including OTX2 in osteogenesis and ZFAND5, MYCN, and RHOX12 in adipogenesis, as experimentally validated for the latter two TFs (**Fig. 2e** and **Supplementary Fig. 2j**). However, differential expression (DE) analysis (**Supplementary Table 3**) revealed that in contrast to CEBPA, PPARG, and RHOX12 cells, MYCN cells lack the expression of *Plin4*, a late stage adipocyte differentiation marker that is crucial for lipid droplet association^45^. This observation is consistent with the smaller and more scattered lipid droplets in MYCN cells compared to CEBPA, PPARG, and RHOX12 cells (**Fig. 2e-f**). While these TFs are all capable of driving MSCs toward the adipocyte lineage, our scTF-seq data suggests that MYCN may do so through a distinct mechanism (as detailed below).

### Characterizing TF dose sensitivity and reprogramming capacity

To quantify the relative transcriptomic change in each cell in function of TF dose (**Fig. 1e**), we projected all cells linked to the 384 TFs into a unified high-dimensional principal component analysis (PCA) space to compare transcriptome variation between each cell and the centroid of control cells (**Fig. 3a** and **Methods**). As expected, the relative transcriptomic alterations were overall greater in TF cells compared to control ones, as well as in functional TF cells relative to their non-functional counterparts (**Supplementary Fig. 3a, b**). By subsequently visualizing transcriptomic change over TF dose (**Fig. 3b**), we found that TFs vary substantially in how their effect is modulated by dose. For example, we observed that some TFs are capable of inducing substantial transcriptomic changes already at a low dose with their effect plateauing at a higher dose, while other TFs were insensitive to dose variation.

**Fig. 3:**
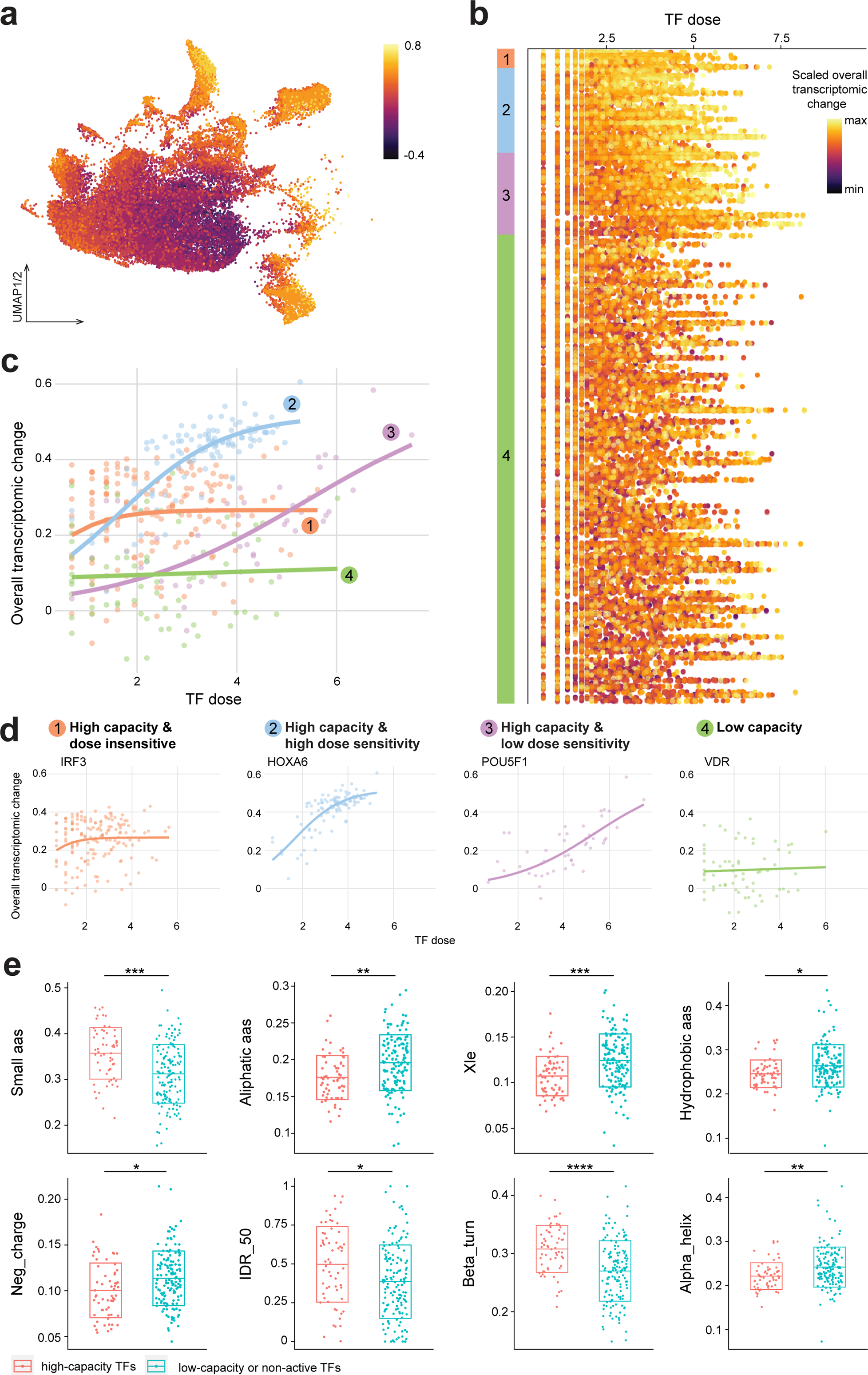
Characterizing TF dose sensitivity and reprogramming capacity. **a** UMAP of the TF atlas colored by overall transcriptomic changes (see **Methods**). **b** Dot plot showing the scaled, overall transcriptomic change of TF-overexpressing cells over TF dose. Each dot represents a cell. Each row represents a TF. Color bars on the left represent TF groups categorized according to dose sensitivity and reprogramming capacity (see color legend in **d**). **c** Scatter plot showing examples of the overall transcriptomic change across TF dose colored for the four TF categories (see legend in **d**). The lines represent the fitted logistic regression. **d** Scatter plot showing the overall transcriptomic change of one representative TF of each TF category across TF dose. The lines represent the fitted logistic regression. **e** Box plots showing the distribution of various TF features (small aas, small amino acids; Aliphatic aas, aliphatic amino acids; Xle, fraction of leucine and isoleucine; Hydrophobic aas, fraction of hydrophobic amino acids; Neg_charge, fraction of negatively charged amino acids; IDR_50, fraction of intrinsically disordered regions; Beta_turn, fraction of beta turns; Alpha_helix, fraction of alpha helices) across high-capacity TFs and low-capacity or non-active TFs. * *P* value ≤ 0.05, ** *P* value ≤ 0.01, *** *P* value ≤ 0.001, **** *P* value ≤ 0.0001, Wilcoxon Rank Sum test followed by FDR correction. See also **Supplementary Fig. 3**.

To better capture TF dose-response relationships, we employed a logistic model (**Supplementary Fig. 3c**) to fit transcriptomic change in function of TF dose. Within this model, we estimated three parameters: Asym, Xmid, and Scal, representing the asymptote, the TF dose at the point of inflection, and the inverse slope at the point of inflection, respectively. As the dose of certain TFs might not have been sufficient to reach saturation, their asymptote sometimes largely surpassed the maximum observed transcriptomic change. To limit potential overestimation, we employed model-inferred transcriptomic changes at the maximum observed dose, defining this property as “TF capacity”. For consistency and comparability across various TFs, we first filtered TFs based on the number of cells and the dose range (**Supplementary Fig. 3d** and **Methods**). 35 TFs were deemed inactive due to their limited number and proportion of functional cells (**Supplementary Fig. 3d** and **Methods**). We then applied the logistic model to a total of 235 TFs of which 11 were finally still excluded due to convergence failures. Leveraging the parameters extracted from the model, we broadly classified the remaining 224 TFs into two main groups: high-capacity and low-capacity ones. To explore the functional relevance of this classification, we analyzed mutational constraint data from gnomAD^46,47^ for human orthologs of these 224 TFs and non-active TFs (if present). We found that genes coding for high-capacity TFs are significantly enriched within the group of genes that are intolerant to loss-of-function mutations (Fisher’s exact *P* < 0.05) (**Supplementary Fig. 3e**). This appears consistent with the notion that these high-capacity TFs may have a more significant impact on cellular and ultimately organismal phenotypes compared to low-capacity ones^46,48^. Within the high-capacity group, we then evaluated whether the effect is dose-dependent and if so, whether this dose sensitivity is either low or high, as defined by the induced transcriptomic response at an average TF dose. This resulted in the categorization of all TFs into four distinct groups (**Fig. 3b-d, Supplementary Fig. 3d** and **Supplementary Table 4**): 1) 8 high-capacity TFs with minimal to no dose sensitivity in the observed dose-range, suggesting an ON/OFF type of mechanism as exemplified by IRF3; 2) 28 high-capacity and high dose-sensitive TFs including HOX and CDX TFs; 3) 28 high-capacity and low dose-sensitive TFs which require a high dose to reach high capacity. The latter includes OCT3/4, a Yamanaka factor playing crucial roles in regulating pluripotency and differentiation (**Fig. 3d**), and whose cellular reprogramming ability has been shown to scale with its expression levels^49,50^; 4) 160 low-capacity TFs that induced no to only very mild transcriptomic effects across a wide dose range.

We next inquired into TF protein features that could explain differences in reprogramming capacity (**Supplementary Table 5** and **Methods**). We found that high-capacity TFs are enriched for features such as small amino acids like proline and serine, intrinsically disordered regions (IDRs), and have a significantly greater propensity to form beta turns which represent energetically favored nucleation points^51^, while being depleted in aliphatic amino acids (including leucine and isoleucine), hydrophobic amino acids, negative charge, and alpha helices (**Fig. 3e** and **Supplementary Fig. 3f**). Similar compositional biases have been revealed as evolutionarily conserved patterns associated with phase-separating proteins including specific TFs and co-regulators whose condensate formation ability is thought to play a key role in gene regulation^52–54^.

### Disentangling the origins of reprogramming heterogeneity: dose and stochasticity

For most high-capacity TFs, the dose is a strong contributor to the observed reprogramming heterogeneity. As seen in **Fig. 3b**, the overall transcriptomic changes of cells varied along doses of high-capacity TFs. However, those changes are summarized into one singular value, lacking specificity regarding gene-specific effects. Therefore, we next investigated whether individual or sets of genes display consistent or different responses to varying doses of the same TF and whether diverse response patterns of genes could facilitate the emergence of different forms of reprogramming heterogeneity. We observed that heterogeneous cell states within a single lineage can be explained by monotonic effects that TF dose has on early and late differentiation genes. For example, the adipogenic gene expression signature (termed adiposcore hereafter) of CEBPA cells strongly correlates with *Cebpa*’s dose (**Fig. 4a**). Indeed, the early adipogenesis regulator *Cebpd* was downregulated, while the master adipocyte differentiation regulator *Pparg* and mature adipocyte markers such as *Fabp5* and *C3* were generally upregulated with increasing doses of *Cebpa* (**Fig. 4b**).

**Fig. 4:**
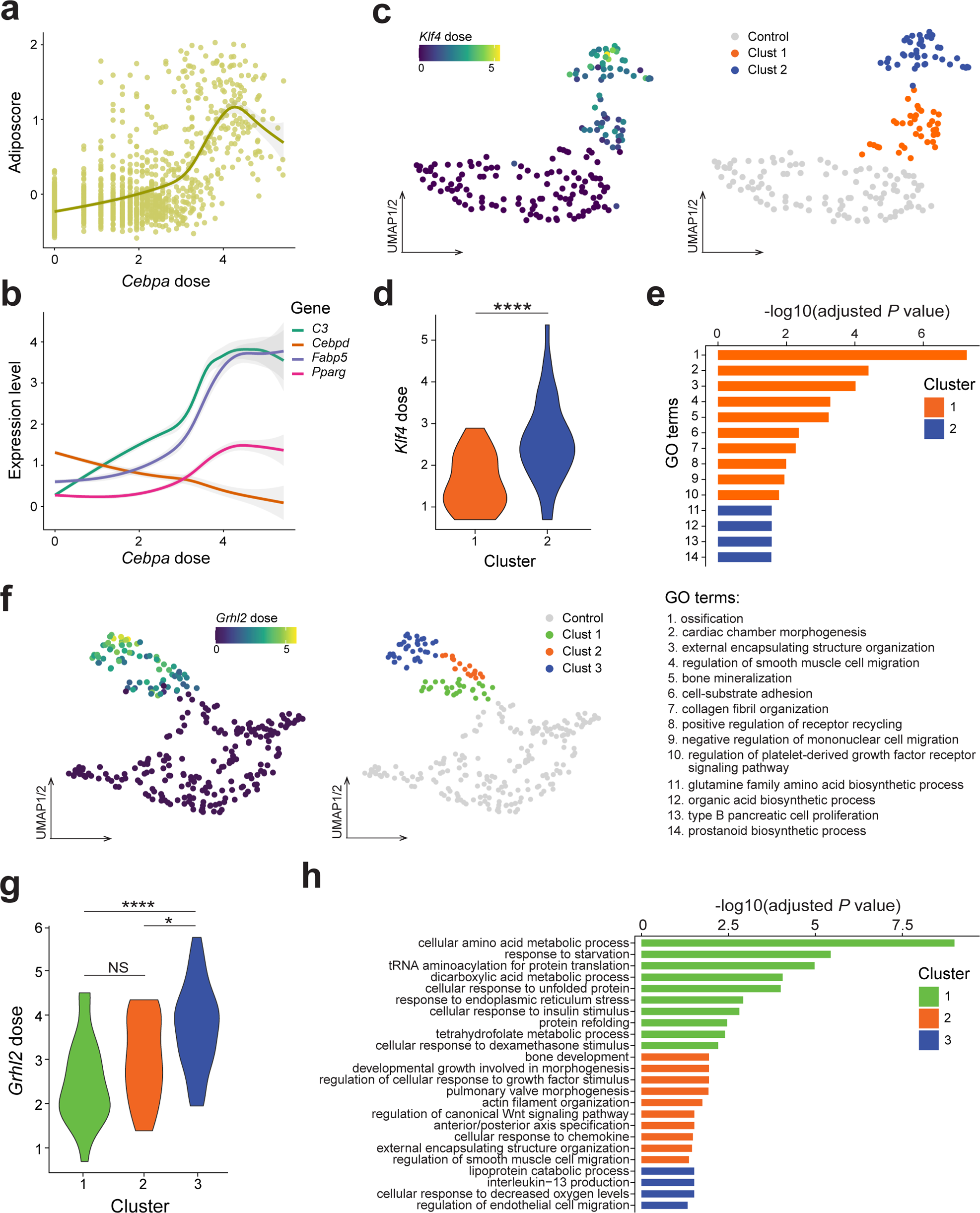
Reprogramming heterogeneity induced by TFs. **a, b** The adiposcore (**a**) and expression level of individual adipogenesis-related genes (**b**) in cells overexpressing *Cebpa* at different doses. The lines represent the fitted loess regression. The bands represent the 95% confidence interval. **c** UMAPs of KLF4 and its batch-paired control cells colored by *Klf4* dose (left) or clusters (right) identified by unsupervised clustering. The clusters containing mainly control cells were merged and labeled as the Control cluster. Clusters containing mainly KLF4 cells are highlighted in color. **d** Violin plot showing the distribution of *Klf4* dose in cells from KLF4 cluster 1 and 2 shown in **c**. **** *P* value < 0.0001, two-sided *t*-test. **e** Top 10 unique GO Biological Process terms identified by GO enrichment analysis on the significantly differentially expressed genes of each KLF4 cluster (shown in **c**). **f** UMAP of GRHL2 and its batch-paired control cells colored by *Grhl2* dose (left) or clusters (right) identified by unsupervised clustering. The clusters containing mainly control cells were merged and labeled as the Control cluster. Clusters containing mainly GRHL2 cells are highlighted in color. **g** Violin plot showing the distribution of *Grhl2* dose among the three GRHL2 clusters shown in **f**. * *P* value < 0.05; **** *P* value < 0.0001; NS, not significant; pairwise two-sided *t*-test followed by FDR correction. **h** Top 10 unique GO Biological Process terms identified by GO enrichment analysis on the significantly differentially expressed genes of each GRHL2 cluster (shown in **f**). See also **Supplementary Figs. 4.1 and 4.2**.

However, we also observed a more complex form of reprogramming heterogeneity where variation in TF dose results in a non-monotonic expression response of genes that drive distinct cell fate specifications. For example, two subgroups that significantly differ in *Klf4* dose and gene expression pattern were uncovered by using clustering analysis (**Fig. 4c, d**; **Supplementary Fig. 4.1a-c**; **Methods**). Genes belonging to the gene ontology (GO) terms such as ossification, cardiac chamber morphogenesis, and regulation of smooth muscle cell migration were significantly upregulated by *Klf4* only at low doses. Conversely, distinct sets of genes associated to various biosynthetic processes were uniquely upregulated in cells highly expressing *Klf4* (**Fig. 4e** and **Supplementary Fig. 4.1d**). These findings suggest that dose variation may direct KLF4 cells to different functional branches regulating differentiation and metabolism^55^, respectively. Besides KLF4, several other TFs exhibited a comparable capacity, including EGR1, ESR2, ETV1, and RUNX2, all of which were able to induce distinct cell states in a dose-dependent manner (**Supplementary Figs. 4.1e, f and 4.2a-c**).

While we could attribute dose as a determining factor in cell fate branching, we uncovered some TFs, including GRHL2, MEIS2, and MYOG, for which cells could be stratified over distinct branches despite featuring a similar dose of the respective TF (**Fig. 4f-h**, **Supplementary Figs. 4.1c and 4.2d-h**). Focusing on GRHL2 as an example, unsupervised clustering stratified GRHL2 cells in three subgroups with cluster 1 showing significantly higher *Grhl2* levels compared to clusters 2 and 3, which exhibited comparable *Grhl2* expression levels (**Fig. 4f, g**). To study the branching properties in greater detail, we computationally inferred trajectories on GRHL2 and batch-paired control cells, using control cells as roots (**Supplementary Fig. 4.2h**). DE analysis among the three GRHL2 clusters along the resulting pseudotime clearly revealed three distinct gene expression profiles that are enriched for distinct cellular processes including lipoprotein catabolic process and regulation of endothelial cell migration (cluster 1), cellular amino acid metabolic processes and cellular responses to insulin stimulus or starvation (cluster 2), and bone development (cluster 3), respectively (**Fig. 4h** and **Supplementary Table 6**). Thus, our findings clearly point to the involvement of factors other than dose that also play an important role in controlling cell fate decisions.

### Dissecting TF-cell cycle interactions and their coordinated impact on reprogramming

One factor that also could cause transcriptomic heterogeneity is the cell cycle given that it is fundamentally associated with the self-renewal and lineage determination of stem cells^28,56^. Yet, our understanding of how the cell cycle interacts with TFs and TF dose in particular, and to what extent it contributes to reprogramming heterogeneity, remains limited. To address this, we leveraged our scTF-seq data to systematically study interactions between TFs and the cell cycle. We first assessed the impact of TF overexpression on cell cycle dynamics. To identify cell cycle phase, each cell was assigned two scores based on its expression of canonical S and G2M markers and further classified into each phase following the default (0) or adjusted (0.1) cutoff on the S and G2M scores (**Supplementary Fig. 1f**, **Fig. 5a** and **Methods**). The latter was applied to rectify the overclassification of ambiguous cells into S/G2M, as observed for example for YAP1, ATF3, and control cells with S and G2M scores between 0 and 0.1 (**Fig. 5a, b** and **Supplementary Figs. 1f, 5a-c**). The proportion of cells in each adjusted phase was compared among all TFs. As expected, the known cell cycle-driving TFs, such as E2F2^57^, T^58^, and MYCN^59^ had the highest proportion of S and G2M cells (**Fig. 5b**). To go further than the discrete phase categorization which overlooks the circular and continuous nature of the cell cycle, we compared the density distributions of cell cycle scores between TF and confluent control cells. The one-dimensional (1D) distributions of S and G2M scores allowed us to specify that E2F2 shifted cells towards high S scores, while T and MYCN increased both the S and G2M scores (**Fig. 5c**). However, the interdependence of S and G2M scores, reflected by the significant correlation of the two scores in control cells, could introduce bias in the interpretation (**Supplementary Fig. 5c**). To address this, we quantified cell density throughout cell cycle progression (**Supplementary Fig. 5d** and **Methods**). Although the two-dimensional (2D) density estimation may have lower sensitivity due to limited numbers of cells, it reflected the trends of cell cycle progression observed in the 1D distributions and clarified that *E2f2* overexpression may not only direct cells into S phase but also block cells in the S phase with very few cells progressing to G2M (**Supplementary Fig. 5e-g** and **Fig. 5c**). This aligned with previous findings showing that stabilized E2F2 activity throughout the cell cycle accelerates G1/S transition in the short term (within 2 days) but initiates replication stress, DNA damage, and apoptosis, thereby impairing long-term cell fitness^60^.

**Fig. 5:**
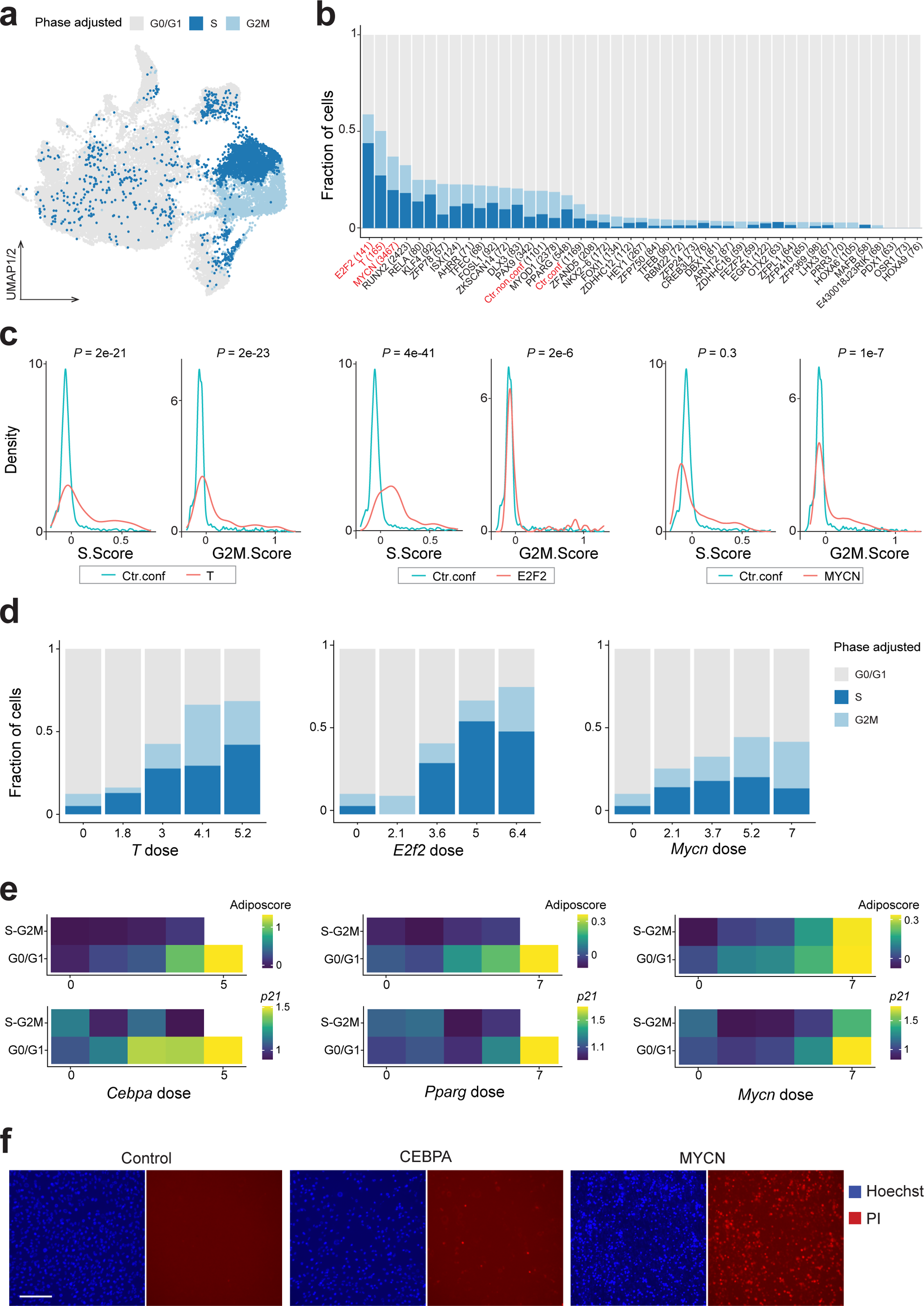
Interactions between TFs, the cell cycle and differentiation (adipogenesis) **a** UMAP of the TF atlas colored by adjusted cell cycle phase (see **Methods**). **b** Bar plot showing the fraction of cells in the adjusted phase for each TF. The total number of cells is indicated in brackets. A Fisher’s exact test was performed between confluent control cells (Ctr.conf) and each TF. Only TFs and the non-confluent control (Ctr.non.conf) that tested significantly (FDR-adjusted p-value < 0.05) are visualized here. **c** Density plots showing the distributions of S and G2M scores of TF cells (T, E2F2, or MYCN in red) compared to control cells (Ctr.conf in green). The adjusted *P* value was generated by the Wilcoxon Rank Sum test followed by FDR correction. **d** Bar plots showing the fraction of cells in each adjusted cell cycle phase across binned doses of *T*, *E2f2*, or *Mycn*. **e** Heatmaps showing the mean expression levels of adipocyte marker genes (adiposcore), and *p21* in CEBPA, PPARG and MYCN cells, which are binned according to their adjusted cell cycle phase and TF dose. Bins with less than 3 cells were excluded (white square). **f** Representative fluorescent images showing the viability of control, CEBPA, and MYCN cells, indicated by propidium iodide (PI) staining in red. Nuclei were stained by Hoechst in blue. Scale bar, 200 μm. See also **Supplementary Fig. 5**.

It is also noteworthy that the proportion of S and G2M cells was generally enhanced by larger *T* and *E2f2* doses, while other TFs including MYCN, RUNX2, and PAX9 exhibited a non-monotonic trend, with the highest fraction of S and G2M cells at middle dose (**Fig. 5d** and **Supplementary Fig. 5h**). Given this observed dose-dependent interactions between TFs and the cell cycle, we next investigated how TFs coordinate cell cycle dynamics and lineage differentiation in the context of TF dose. MYCN is of particular interest here given that we found different patterns of cell proliferation and differentiation interactions in MYCN cells compared to well-known adipogenic TFs. For instance, cell proliferation and the adiposcore were mutually exclusive in cells overexpressing *Cebpa* and *Pparg* (**Fig. 5e**), consistent with the notion that lineage differentiation including adipogenesis requires cells to exit the cell cycle^28,56,61^. Indeed, *p21*, encoding a cyclin-dependent kinase inhibitor exploited to ensure a harmonized transition between permanent cell cycle exit and adipocyte differentiation^61^, was upregulated at high doses of *Cebpa* and *Pparg* (**Fig. 5e**). However, the cell cycle exit and cell differentiation were decoupled in high dose MYCN cells, as evidenced by the high adiposcore and *p21* expression in S and G2M MYCN cells and the observed concomitant of lipid droplet accumulation and increasing nuclei counts for MYCN cells (**Fig. 2e** and **Supplementary Fig. 2j**). This aberrant differentiation in the high *Mycn* context likely comes at a cost given that evident cell death was observed in this condition (**Fig. 5f**). These collective results underline the intricate interplay between TFs, TF dose, cell cycle dynamics and lineage differentiation.

### Synergistic or antagonistic effects of TF pairs depend on the dose

Transcription factors do not operate in isolation^62^ and the dose of a TF in a specific cell should in principle always be considered relative to that of another TF^63^. Yet, how the dose of one TF impacts the effects of another TF is barely understood due to the complexity underlying combinatorial analysis. To start addressing this, we selected a subset of key TFs that tended to have the strongest lineage differentiation potential in our scTF-seq screen of individual TFs, including CEBPA, PPARG and MYCN for adipogenesis, MYOG for myogenesis, and RUNX2 for osteogenesis. Subsequently, we performed combinatorial scTF-seq experiments for these TFs (**Fig. 6a** and **Methods**).

**Fig. 6:**
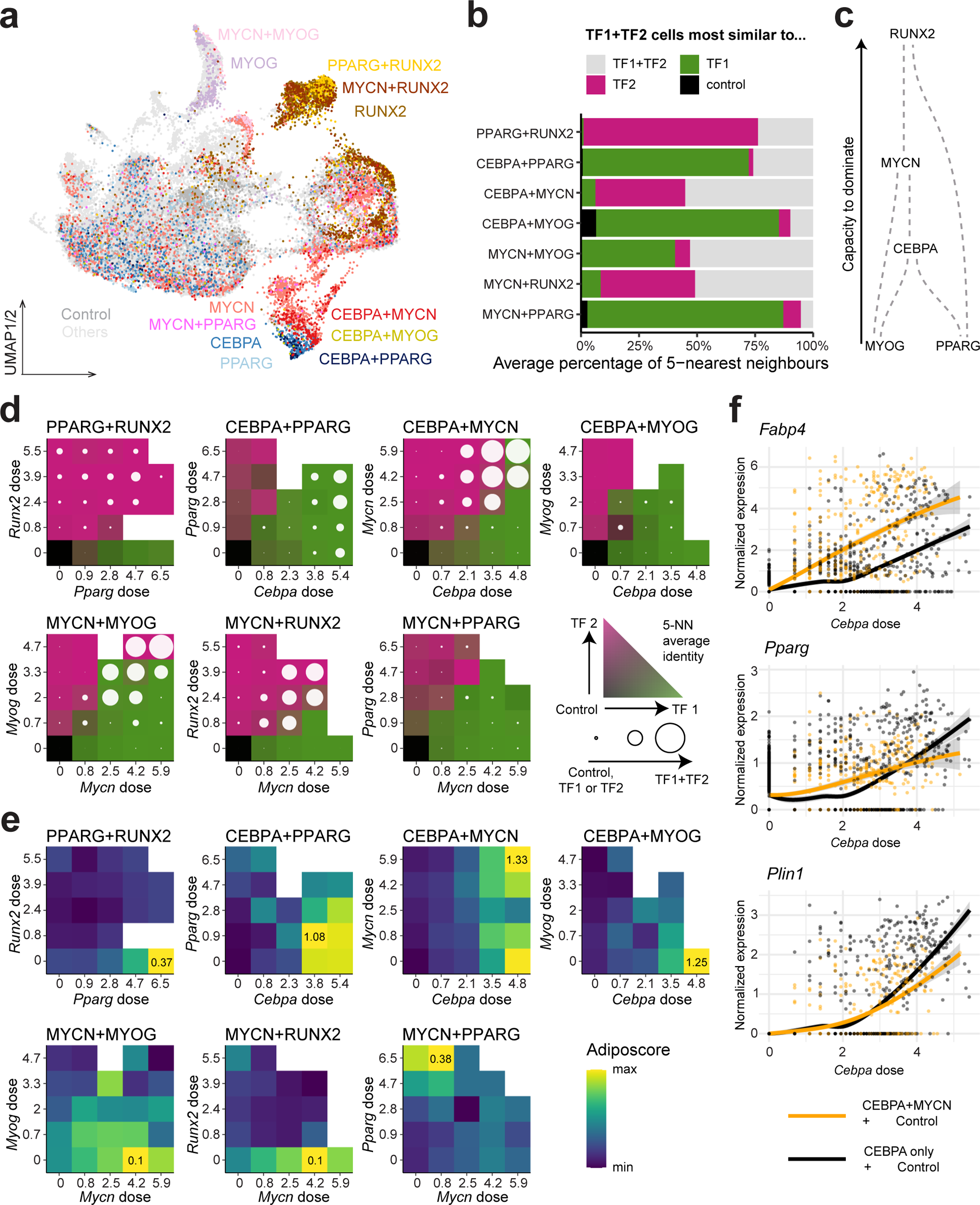
Dose-dependent effects of interactions between TFs. **a** UMAP representation showing the positions of single- and combinations of TFs with respect to all other tested TFs and control cells. **b** Percentage of 5-nearest neighbors to which each combinatorial TF (TF1+TF2) cell is closest. **c** Schematic representation of the dominance of each TF inferred from panel (**b**). For example, RUNX2 dominates over other TFs because when combining RUNX2 with another TF, the combination cells are transcriptomically the most similar to RUNX2 cells. **d** Percentage of 5-nearest neighbors to which each combination cell is closest within a pair of dose bins. Bins were determined based on uniformly splitting the interval between 0 and the maximal dosage, with an additional bin at a dose of 0. The color scale represents the percentage of cells closest to TF1, TF2 or the control cells respectively, represented using a bilinear interpolation between green, pink, and gray. The circle represents the percentage of cells closest to the TF1+TF2 cells, with a full circle meaning that all cells were closest to TF1+TF2. **e** Transcriptomic adiposcore between different dose bins. The color scale ranges from the maximal and minimal adiposcore for each combination of TFs. **f** Dose response-curves for control cells with only CEBPA cells (black), compared to those for control with MYCN*+*CEBPA cells (orange). The band represents the 95% confidence interval on the smoothened mean using locally estimated scatterplot smoothing. See also **Supplementary Fig. 6**.

Utilizing the single-cell readout, we assessed whether a TF pair (TF1+TF2) could induce a cell state different from that observed in cells overexpressing only one of the two TFs. To do so, we quantified whether a given cell with a combinatorial identity would be more similar to cells of the same identity or to those of the individual TF’s identities. We found that the transcriptomic response of one TF dominates (**Fig. 6b**) even across combinations, enabling the construction of a non-cyclic graph of TF dominance (**Fig. 6c**). Yet, the combination of TFs also created a unique transcriptomic profile as observed for MYCN+RUNX2, MYCN+MYOG, and CEBPA+MYCN (**Fig. 6b**). To some extent, these unique states were hybrids of the two original single TF states, given that many genes were differentially expressed only compared to one of the single TFs (**Supplementary Fig. 6a**). Nonetheless, these hybrid states were accompanied by the significant up- and downregulation of a distinct set of genes (**Supplementary Fig. 6a**). Among the uniquely upregulated genes of CEBPA+MYCN were several adipogenesis-related genes (*Fabp4* and *Gpd1l*), indicating a synergistic effect of CEBPA and MYCN.

Since all screened TF pairs contain at least one adipogenic TF, we then assessed their combinatorial effects on adipogenic capacity by quantifying the transcriptomic adiposcore. We observed that TFs targeting both the adipogenic or different lineages have respectively a strongly synergistic or antagonistic effect on adipogenic capacity (**Supplementary Fig. 6b**). These findings were substantiated by the respectively higher or lower lipid score for MYCN+CEBPA or MYOG+CEBPA compared to CEBPA cells (**Supplementary Fig. 6c, d**).

We next investigated combinatorial effects of two TFs in the context of TF dose. We observed that dominance of one TF over another can be strongly influenced by the relative dose of the two TFs. For example, any TF with a much greater dose than another TF was able to overcome the dominant effect of this other TF, with the exception of PPARG due to its low reprogramming capacity (**Fig. 6d**, **Supplementary Fig. 6e** and **Supplementary Table 4**). Furthermore, a high dose was typically required to produce the unique combinatorial cellular state (**Fig. 6d**). Also, the dose sensitivities of some TFs changed when put into competition. For example, MYOG was defined as a high-capacity and dose-insensitive TF, whereas it was mostly dominated by other high-capacity and dose-sensitive TFs at low doses (**Fig. 6d**, **Supplementary Fig. 6e** and **Supplementary Table 4**). Despite having a lower dose sensitivity than CEBPA, MYCN dominated over CEBPA when they were present at a similar dose (**Fig. 6d**, **Supplementary Fig. 6e** and **Supplementary Table 4**). Next to the overall cell identity, the synergistic or antagonistic effects of TF pairs on adipogenic capacity were also dependent on TF dose (**Fig. 6e**). Interestingly, we found that interactions between TFs can be non-monotonic to dose. For instance, although the interaction between CEBPA and MYCN resulted globally in a synergistic effect (**Supplementary Fig. 6b**), MYCN at an intermediate dose even antagonized adipogenic capacity in high dose CEBPA cells (**Fig. 6e**). Conversely, the highest adipogenic capacity of the CEBPA+PPARG combination was observed at a low *Pparg* dose, a rather surprising finding given PPARg’s well-known role as master regulator of adipogenesis^43^.

Finally, we identified that the dose-dependent synergism of TF pairs can be gene-specific. As exemplified by CEBPA+MYCN, some key adipocyte markers, such as *Fabp4* and *Adipoq*, were synergistically regulated, while other adipogenesis-related genes, such as *Pparg* and *Plin1*, switched between a synergistic and antagonistic relationship along *Cebpa* dose (**Fig. 6f** and **Supplementary Fig. 6f**). The fact that these interactions are often non-linear and gene-specific, reflects a complex regulatory architecture which requires a dose-specific transcriptomics approach to be fully understood.

## Discussion

Numerous studies have emphasized the transformative impact of adjusting the TF dose on molecular and cellular states^9,13,15,64–67^. The significant degree of cellular heterogeneity typically observed upon TF overexpression in *ex vivo* experiments contrasts however with the precise control of cell fate alterations observed *in vivo*. This discrepancy signals a gap in our understanding of how cellular programs intricately respond to TF dose variation. To address this gap, we developed scTF-seq, demonstrating its versatility in enabling i) the identification of experimentally validated lineage regulators and associated functional modules, rendering the resulting scTF-seq atlas a comprehensive reference for identifying TFs that induce phenotypes of interest (**Fig. 2**); ii) the systematic and quantitative mapping of the influence of TF dose on cell reprogramming at the single-cell level, which constitutes a unique feature compared to other large-scale single-cell^16,18–20,68^ or bulk^3,69,70^ TF screening strategies (**Figs. 3-6**).

By exploring this intricate relationship between TF dose and function, we were able to stratify TFs into four distinct categories: low versus high-capacity TFs with the latter further classified as ‘not,’ ‘low,’ or ‘high’ dose-sensitive (**Fig. 3**). While the biological meaning of these TF groups remains unclear, one possibility is that the capacity reflects the extent of the respective TF’s functional impact. This notion is supported by the higher intolerance to loss-of-function mutations present in human orthologs of high-capacity TFs than low-capacity or non-active TFs^46,48^. The dose sensitivity may also be highly relevant to how TFs exert their function in response to stimuli or developmental signals. For example, IRF3 is a high-capacity TF consistent with its role as a key driver of virus-induced type I interferon production. However, only a small fraction of the IRF3 protein pool needs activation to elicit an antiviral response, while the activation of the IRF3 pathway is controlled by a ‘licensing’ mechanism to prevent excessive production of interferons^32,71^. This phenomenon consists with the characterization that IRF3 is not dose-sensitive. Furthermore, many *Hox* and *Cdx* TF family members feature a high capacity and high dose sensitivity, aligned with their known influence in development through a concentration gradient^72,73^. In contrast, OCT3/4 is a high-capacity, but low dose-sensitive TF, which may be consistent with observations that highest reprogramming efficiencies tend to be reached at highest *Oct3/4* overexpression levels^49,50,66^. A large proportion of TFs exhibited a low capacity though, exemplified by vitamin D3 receptor VDR, which is likely ineffective without sufficient supply of its ligand. We thus cannot exclude the possibility that the classification of some of the TFs would change depending on the probed system, stimuli or even utilized approach or model. Despite this possible classification variation, we uncovered a clear enrichment among high-capacity TFs of features linked to phase-separating properties^51,52,54^. This observation hints at a potentially compelling connection between reprogramming capacity and condensate formation in cellular processes.

Within the realm of high-capacity TFs, our findings illuminate the significant role that dose plays in introducing reprogramming heterogeneity (**Fig. 4**). This was for example manifested in diverse gene response patterns resulting in dose-dependent cell fate branching. Nevertheless, not all observed branching events could be solely attributed to dose, as shown for GRHL2. Its target cells were stratified into three distinct clusters with two featuring a comparable dose. The molecular mechanisms behind this dose-independent, potentially stochastic cell fate branching remain poorly understood. It may reflect the stochastic nature of gene transcription, arising from the dynamic interplay among transcriptional processes (such as TF-DNA binding kinetics), epigenetic modifications, and post-transcriptional events in individual cells^74–78^. Alternatively, these mechanisms may be influenced by more deterministic factors. The latter may include the cell cycle phase during initial TF overexpression^12^, although our observations indicate that the influence of the cell cycle can also extend beyond the starting cell population (**Fig. 5**). Indeed, several TFs, including master regulators like RUNX2 and PAX9, exhibit complex, non-monotonic patterns indicative of an interplay between the cell cycle and TF dose. This implies that such TFs can function as rheostats, regulating dose-dependent entry into the cell cycle to control terminal differentiation, consistent with previous observations for the TF MITF^79^. Such variability in cell fate branching holds biological significance, emphasizing the need to systematically catalog distinct dose-response patterns to advance our understanding of the molecular mechanisms that govern these intricate cellular transitions.

Our study also pointed to certain TFs that challenge the conventional requirement for cell cycle exit in terminal differentiation. For instance, MYCN, capable of triggering adipogenesis, displayed a unique dynamic where overexpressing cells actively cycled while concurrently expressing adipogenic genes in a dose-dependent manner (**Fig. 5**). Unraveling how MYCN regulates this intriguing, likely rare state will necessitate more investigation, but it possibly reflects MYCN’s powerful, pleiotropic role in controlling a wide range of cellular processes underlying organogenesis^59^.

Moreover, our study underscores the critical role of TF dose when overexpressing multiple TFs to enhance reprogramming effects, as highlighted by the instances where relationships between TFs shifted from antagonistic to synergistic based on dose (**Fig. 6**). This frequently non-linear and gene-specific nature of TF interactions may reflect the diverse roles of implicated TFs in mediating various aspects of gene regulation, such as controlling chromatin accessibility, regulatory element interactions, and gene activation^15,80^. The observed complexity in TF interactions points to the significant challenge of determining optimal dose regimes for sets of TFs required to generate specific cell states.

In summary, our study not only sheds light on the pivotal role of TF dose in cellular reprogramming but also opens avenues for further exploration. scTF-seq’s agnostic nature to the cell system or species coupled with its potential to provide insights into the regulatory properties of TFs in various contexts positions this assay as a valuable asset for future research. However, certain limitations of the current study should also be acknowledged such as the current lack of temporal resolution in the scTF-seq atlas, emphasizing the need for investigating reprogramming across multiple time points. Additionally, future iterations of the analysis should consider incorporating additional data modalities, such as chromatin accessibility, to unravel the molecular mechanisms underlying observed TF dose effects. This integrative approach would hold promise for deepening our understanding of TF-mediated changes in the chromatin landscape and their implications for cellular reprogramming.

## Methods

### Experimental model and subject details

#### Cell culture and differentiation

Both HEK293T (ATCC, no. SD-3515) and C3H10T1/2 (ATCC, no. CCL-226) cells were maintained in basic culture medium containing high-glucose DMEM with GlutaMax and pyruvate (Thermo Fisher Scientific, no. 10569010), 10% FBS (Gibco, no. 10270106) and 1x Penicillin-Streptomycin (Life Technologies, no. 15140-122). All C3H10T1/2 cells were maintained below passage 15.

For *in vitro* adipogenic differentiation, C3H10T1/2 cells were first cultured in the basic culture medium supplemented with 100 ng/mL BMP4 (RnD systems, no. 314-BP-010) for 3 days. Then the induction medium was added for 2 days, which was composed of the basic culture medium and MDI cocktail containing 1 µM dexamethasone (Sigma-Aldrich, no. D4902-25MG), 0.5 mM 3-isobutyl-1-methylxanthine (Sigma-Aldrich, no. I5879-5g) and 167 nM insulin (Sigma-Aldrich, no. I9278-5ml). The cells were maintained in the basic culture medium supplemented with 167 nM insulin until harvesting.

All cells were placed at 37 °C and 5% CO2 in a humidified incubator. Prior to use, cells were washed with PBS (Thermo Fisher Scientific, no. 14190169), dissociated with Trypsin-EDTA (Thermo Fisher Scientific, no. 25200056; 0.05% for HEK293T and 0.25% for C3H10T1/2), resuspended with basic culture medium, filtered using 40 µm strainers and counted with Trypan blue live-dead stain (Thermo Fisher Scientific, no. T10282) using a Countess automated cell counter (Invitrogen, no. AMQAX2000).

#### Lentivirus production

Lentiviral packing was performed using lipofectamine 2000 according to the manufacturer’s instructions. First, 10 µL of Opti-MEM (Thermo Fisher Scientific, no. 31985070) and 0.375 µL of lipofectamine 2000 Transfection Reagent (Thermo Fisher Scientific, no. 11668027) were thoroughly mixed. A mix of 0.075 µg lentiviral expression plasmid containing the individual TF ORF (or mCherry as control) and TF-ID of interest, 0.075 µg 3^rd^ generation lentivirus packaging plasmid mix (pRSV-Rev:pMDLg/pRRE:pCMV-VSV-G=1:1:1) and 10 µL of Opti-MEM were prepared. Then, two mixes were added together and incubated for 30 min at room temperature. HEK293T cells were cultured and prepared as described above, and seeded in the individual well of 96-well plates at ∼95% confluency. After incubation, the transfection mix was added to freshly seeded cells. Medium was changed 12 h after transfection. 48 h post transfection, the supernatant containing virus particles was harvested, and dead cells were removed by centrifugation at 300g for 5 min. As a control, pBOB-GFP (kindly provided by Dr. Jiahuai Han at Xiamen University) plasmid with constitutive GFP expression was transfected.

#### Lentivirus transduction

C3H10T1/2 cells were seeded 12 h prior to transduction at 10-20% density. Transduction medium was prepared by mixing the supernatant containing lentivirus particles and the basic culture medium in 1:1 ratio, supplemented with polybrene (Sigma-Aldrich, no. TR-1003-G) at a final concentration of 10 µg/mL. Plated cells were then treated with the transduction medium and centrifuged at 1300 g for 30 min at 37 °C. Medium was refreshed after 24 h. After 48 h, cells were selected using 2 µg/mL Puromycin (Thermo Fisher Scientific, no. A1113803) during 48-72 h. Puromycin-resistant cells were cultured in the basic culture medium to recover for 24 to 48 h.

### Experimental details

#### Oligos

Primer sequences used in this study can be found in **Supplementary Table 7**.

#### Barcoding and cloning of TF ORF libraries

Barcoded doxycycline-inducible lentiviral expression vectors carrying TF ORFs (pEXPRESS) were generated individually using the Gateway cloning system in two steps. In the first step, barcoded destination vectors were generated by introducing random nucleotides to the upstream region of the 3’ LTR of pSIN-TRE-GW-3xHA-puroR vector (kindly provided by Dr. Didier Trono at the EPFL). Two fragments were amplified from the pSIN-TRE-GW-3xHA-puroR vector using Kapa HiFi ready mix (1X; Roche, no. 07958935001) with 0.3 µM Enrich_F3 and 0.3 µM pTREP-BC-RamR), 0.3 µM pTREP-vec-R and 0.3 µM pTREP-BC-RamF, respectively, following the program: 1) 98 °C for 3 min, 2) 98 °C for 30 s, 3) 63 °C for 30 s, 4) 72 °C for 5 min, repeat step 2-4 for 15 cycles, 5) 72 °C for 5 min. After purifying both PCR products using a 1% agarose gel and a gel purification kit (Zymo, no. D4007), the two fragments were assembled using a Gibson assembly mix (NEB, no. E2611S) according to the manufacturer’s instructions. Assembled plasmids (termed pTREP-ID vector hereafter) were then purified using a DNA Clean and Concentrator purification kit (Zymo, no. D4014) and transformed into One Shot ccdB Survival 2 T1R resistant competent cells (Thermo Fisher, no. A10460). Successful colonies were then inoculated to growth medium (Luria Broth) containing Ampicillin (AppliChem, no. A0839) and Chloramphenicol (Brunschwig, no. ACR22792-0250) for miniprep (Zymo, no. D4015) and validation. In the second step, TF ORFs were transferred from generated entry clones^81^ to pTREP-ID vectors using LR Clonase II enzyme mix, producing pEXPRESS plasmids. Stbl3 one shot competent cells (Invitrogen, no. C737303) were then transformed with pEXPRESS and grown on ampicillin (100 ug/mL) plates overnight. Colonies were picked and transferred to LB with ampicillin for miniprep or midiprep. The barcodes (termed TF-IDs hereafter) and TF ORF on the pEXPRESS were examined by Sanger sequencing with the usage of microsynth standard primers: EGFP-C-Rev and TET-CMV-for.

#### Single TF overexpression screen, 10x scRNA-seq sequencing and TF-ID enrichment

TF-IDs used in each 10x scRNA-seq experiment were checked for similarity prior to use. Only TF-IDs with a hamming distance greater than 2 nts were retained within each experiment for multiplexing purposes. All scRNA-seq experiments were performed using Chromium Single Cell Expression 3’ Reagent Kits (10x genomics; v2 Chemistry for experiments 1-2, no. PN-120237; v3 Chemistry for experiments 3-8; 10x genomics, no. PN-1000268) following the manufacturer’s instructions. C3H10T1/2 cells were transduced with the lentivirus particles carrying each barcoded TF ORF expression vector individually. Puromycin selection was performed to enrich successfully transduced cells. TF expression was induced by Doxycycline (2 µg/mL; Sigma-Aldrich, no. D9891-1G) treatment during 5 days in cells placed in a basic culture medium refreshed every 48 h. Since C3H10T1/2 cells might undergo spontaneous differentiation once reaching 100% confluency, mCherry was over-expressed under the same condition in both non-confluent and confluent C3H10T1/2 cells as a control. Unless specified, all control cells were considered in subsequent analyses by default. After that, cells were harvested as described above, pooled together and loaded in the 10x Genomics Chromium Controller targeting 8,000-10,000 cells per experiment.

To specifically enrich the TF-ID, an additional PCR amplification targeting the 10x barcode, UMI and TF-ID was conducted using the full-length cDNA product of the 10x scRNA-seq library. The cDNA library (6 ng), BC_vec_target_10X_F1 vector-specific forward primer (0.3 µM), Truseq_universal_adaptor (0.3 µM) and Kapa HiFi ready mix (1X) were used following the program: 1) 98 °C for 30s, 2) 10 cycles of 98 °C for 10s, 63 °C for 20s and 72 °C for 30s, 3) 72 °C for 5 min. The resulting amplicons were then purified using Ampure beads (2.5X; Labgene, no. CNGS-0050) and further amplified to generate TF-ID-enriched libraries compatible with 10x cDNA libraries with Truseq_D7_adapter (0.3 µM), Truseq_universal_adapter (0.3 µM) and Kapa HiFi ready mix (1X) following the program: 1) 98 °C for 30s, 2) 4 cycles of 98 °C for 10s, 63 °C for 20s and 72 °C for 30s, 3) 72 °C for 5 min. The TF-ID-enriched libraries were then purified twice using 0.6X Ampure beads and pooled with the regular 10x sequencing libraries, which were then sequenced together on the Illumina NextSeq 500/Hiseq 4000 platform using the dual-index configuration following manufacturer’s instructions to obtain a mean depth of 50,000 reads per cell.

#### TF pair screening

To generate data with combinations of TFs, C3H10T1/2 cells were transduced with the barcoded TF ORF expression vector for the first TF and selected with puromycin according to the previously described procedures. Thereafter, the second TF barcoded with a different TF-ID was transduced into the selected cells (virus MOI around 3) that already contained the expression cassette of the first TF. The overexpression of both TFs was induced by doxycycline following conditions already described above.

#### Validation of the adipogenic capacity of single TFs or TF pairs

By using the previously described conditions, C3H10T1/2 cells were transduced with the barcoded TF ORF expression vector with mCherry (control), individual adipogenic TFs, or TF pairs of interest, followed by the Puromycin selection and 5 days of Doxycycline-induced TF overexpression. Thereafter, cells were fixed with 4% PFA (Electron Microscopy Sciences, no. 15714) for 15 min at room temperature, permeabilized with PBS and triton (0.3%, Applichem, no. A4975.0100) and stained with fluorescence dyes: Bodipy 10 µg/ml (boron-dipyrromethene; Invitrogen, no. D3922) for lipids and DAPI (1:5000; Sigma-Aldrich, no. D9542-1MG) for nuclei. Cells were incubated with the dyes in PBS, for 30 min in the dark, washed twice with PBS, and imaged. Stacks of 10 images per well (96 well plate) were collected for each replicate for both blue and green channels using a 20x/0.8 objective. To accurately estimate and represent differences in adipocyte differentiation, an image pre-processing and quantification algorithm was used following the developer’s instruction^82^. Lipid score was then defined by the thresholded Bodipy area divided by Nuclei counts.

#### Cell death staining

By using the previously described conditions, C3H10T1/2 cells were transduced with mCherry (control), *Cebpa*, and *Mycn* followed by Puromycin selection and 5 days of Doxycycline-induced TF overexpression. After this, cells were stained with fluorescence dyes: Propidium iodide 750 nM (PI; Solarbio, no. CA1630) for cell viability and Hoechst 5 µg/ml (Thermo Fisher Scientific, no. H3570) for nuclei. Cells were incubated with dyes in PBS for 30 min in the dark, washed twice with PBS, and imaged.

### Quantification and statistical analysis

#### 10x scRNA-seq data preprocessing and quality control

Sequencing data were aligned and quantified using Cell Ranger against the GRCm38 (mm10, Ensembl release 96) mouse reference genome with default settings to generate count matrices of genes x cell barcodes. To match TF-IDs to cells, sequencing data were also mapped to the pEXPRESS vector sequence where each nucleotide of the TF-IDs was replaced by “N”. Then, TF-IDs from aligned reads at the location of “Ns” were extracted with 1 nt mismatch allowed and matched to the corresponding cell barcodes and UMIs using an in-house framework that we developed to assure reproducibility: TFseqTools (https://github.com/DeplanckeLab/TFseqTools), which at the end yielded read/UMI matrices containing TF-IDs x cell barcodes.

All the data were then loaded on R (R version 4.1.0). Quality control and TF-ID assignments were performed for each experiment individually. Only cell barcodes detected both in the 10x and TF-ID enrichment libraries and having more than 5 TF-ID reads were retained. Approximately 40% of the cells had reads aligning to more than one TF-ID (**Supplementary Fig. 1d**). To assign a TF to a cell, cells were ordered based on the proportion of their main TF-ID (defined as the percentage of reads aligning to the most abundant TF-ID) from high to low for each experiment. Knee point detection was then applied to find the inflexion point of the curve using the kneepointDetection function of the R package SamSPECTRAL^83^. Cells with a percentage lower than the knee point (on average above 80%) were considered as doublets or heavily contaminated by ambient RNAs and filtered out. Thereafter, the dataset of remaining cells from each experiment was analyzed with Seurat (v3 and v4)^84^. TF dose was computed by using the natural logarithm transformation of one plus UMIs of the assigned TF-ID. Genes detected in fewer than 3 cells were removed. Low quality cells were filtered out using the median absolute deviation from the median total UMI numbers and gene numbers, implemented in the isOutlier function of package SCRAN^93^, using a cutoff of nmads = 4 to 6 of the lower end tail depending on the gene expression matrix of individual experiments. Cells with more than 10% or 15% of mitochondrial gene expression, 40% of ribosomal RNAs and less than 75% of protein-coding genes were also filtered out. TFs having less than 8 cells were excluded.

#### Classification of cell cycle phase

Cell cycle phase was identified per experiment by the CellCycleScoring function from Seurat using a default threshold (0) of S and G2M phase scores. A more stringent threshold (0.1) of phase scores was applied for adjusted phase partitions. When focusing on lineage differentiation, the default phase was used to remove cells that might be preparing for entering the S phase or exiting G2M phase to eliminate the cell cycle confounder. The adjusted phase was employed for selecting cells that are definitely characterized as S or G2M states. The usage of adjusted phase partitions was specified in downstream analyses.

#### Selection of functional cells

Cells of each TF in each experiment were processed per cell cycle phase (default) together with their batch-paired mCherry-overexpressing control cells, which limits confounders of the batch effect and the cell cycle. After normalization and scaling on the 2000 highly variable features as implemented in Seurat, principal component analysis (PCA) was performed with a maximum of 20 principal components (PCs). Only significant PCs that had a *P* value smaller than 0.05 as calculated by Jackstraw were retained for downstream analysis. In the PCA space, derived from the significant PCs, three centroids were delineated for confluent, non-confluent, and all control cells to mitigate biases due to variation among control cells and their confluency. The Euclidean Distance between individual cells and the three designated centroids of the control cells were computed. After assessing a gradient of percentile-based thresholds rather than a uniform fixed value for all TFs, the 80th percentile of the distance of control cells to their respective centroids was employed to identify functional TF cells which have greater distances than the threshold.

#### Data integration and clustering analysis

Data integration was performed following the guidelines of Seurat. TFs that have less than 8 cells or 5 functional cells were excluded. 2,000 highly variable features were retained for integration and data scaling. The dimensionality reduction was performed via PCA with 200 PCs for the integration of all cells or 50 PCs for the integration of control cells and functional TF cells in G1 (default phase) with or without all cells in S/G2M (default phase). The clustree function from clustree package was applied to find an optimal resolution for clustering^86^. The exact resolution of clustering was specified in downstream analyses when applicable. Cells and clusters were visualized using uniform manifold approximation and projection (UMAP).

#### Differential expression and enrichment analyses

Differential expression analysis was performed on all detected genes by using generalized linear models with batch as a covariate using edgeR^87^. Marker genes of cell types of interest and hallmark gene sets were downloaded from MSigDB v2023.1.Mm and PanglaoDB^88^. A clustering resolution 0.2 was used for enrichment analysis of clusters on hallmark gene sets. A customized gene set containing more mature adipocyte markers^89^ (*Fabp4*, *Lpl*, *Pparg*, *Lipe*, *Adipoq*, *Cd36*, *Plin4*, *Plin2*, *Plin1*, *Cebpa*, *Cebpb*, *Cidec*, and *Cidea*) was used to compute the adipocyte module score (referred to as the adiposcore) by using the AddModuleScore function from Seurat. Gene set enrichment analysis was performed using the package fgsea^90^. Due to the relatively small number of gene sets (adipocytes, chondrocytes, myoblasts and osteoblasts) being analyzed, a false discovery rate (FDR) cutoff 5% was used. Gene ontology (GO) enrichment analysis was implemented by using the enrichGO function from clusterProfiler^91^. GO terms with more than 50% genes overlapping were excluded.

#### Cellular similarity analysis

To assess cellular similarities and identify TFs that have similar biological functions (referred to as functional modules), pairwise Pearson correlation coefficients were computed using the rcorr function of the Hmisc package^92^. This was conducted in the PCA space, constructed from the first 50 PCs, and was inclusive of both control and functional TF cells in G1 (default phase).

#### Transcriptomic change, TF dose sensitivity and reprogramming capacity analyses

To quantify the transcriptomic change of the TF cells relative to the control cells, a negative Pearson correlation between each cell and the centroid of control cells was computed in a dimensional space, derived from the top 200 PCs and integrated with all TF and control cells in adjusted G1 phase. The resultant values were then adjusted by subtracting the mean of the negative correlation between control cells and their centroid.

To study and compare the regulatory potential among individual TFs, a self-starting nonlinear least squares logistic model was used to fit the TF-induced transcriptomic changes against the TF dose using the nls function from the package stats (algorithm = ‘SSlogis’). TFs with fewer than 30 cells and low dose variation (i.e. a minimum dose exceeding 2 or a maximum dose falling below 3.8, the latter corresponding to the 15th percentile of the maximal doses of all TFs) were omitted. TFs that have fewer than 5 functional cells or less than 15% functional cells were categorized as inactive. The logistic model estimates three parameters (Asym, Xmid, and Scal) and predicts the average response at every dose. Based on these parameters, the TFs were classified as low/high capacity and no/low/high dose-sensitivity TFs as described below and schematised in **Supplementary Fig. 3d**. High-capacity TFs were defined as TFs with predicted maximum transcriptomic changes equal or above 0.25. A TF was classified as dose-independent if it met the following criteria: Xmid ≤ log1p(2) or Scal ≥ 6 or (Scal > 4 & Xmid equals the largest dose of that TF). For those TFs that were both dose-dependent and exhibited a high capacity, the predicted transcriptomic changes at the average dose of all TFs were used to stratify these TFs into high or low dose sensitivity groups (see **Supplementary Fig. 3d**).

#### TF feature enrichment

TF features including amino acid content, intrinsically disordered region score, and beta turn fraction as listed in **Supplementary Table 5** were calculated by using the phase separation analysis and prediction (PSAP) classifier^52^. A Wilcoxon Rank Sum test followed by FDR correction was applied to compare the distribution of TF features between high-capacity TFs and low-capacity or non-active TFs. An adjusted *P* value < 0.05 was considered statistically significant.

#### Analysis for intolerance to loss of function

The mutational constraints quantified from variation in 141,456 humans were downloaded from gnomAD^46,47^. A Fisher’s exact test was applied to compare the pLI (probability of being loss-of-function intolerant) score and the LOEUF (loss-of-function observed / expected upper bound fraction) across human orthologs of high-capacity TFs and low-capacity or non-active TFs, by using a typical threshold of 0.9 for pLI and a gnomAD-suggested threshold of 0.6 for LOEUF, respectively.

#### Cell fate branching analysis

To identify TF cells that underwent specialized cell fate branching, clustering analysis was performed on control and functional TF cells in G1 (adjusted phase) using a resolution of 0.8 using the function FindCluster from Seurat. Clusters that were predominantly composed of control cells were classified as control clusters. The remaining clusters were annotated as functional clusters. A TF with a certain proportion of its cells (15-85% cells for that TF) in at least two functionally distinct clusters was deemed to be a candidate steering cell fate branching. TFs that had less than 30 cells in total, less than 10 cells and 15% TF cells in each functional cluster were neglected.

To track the cellular state divergence over pseudotime, cells of each TF candidate in G1 (adjusted phase) were pooled with their batch-paired control cells. Monocle3^93,94^ was employed to conduct dimensionality reduction (to 30 or 50 dimensions), batch correction (for TFs enrolled in more than one batch), clustering (resolution 0.1-0.3), and trajectory inference using the paired control cells as roots.

#### Comparisons of cell cycle progression

Since all TF cells were confluent, only confluent mCherry-expressing cells were used as the control in this analysis to avoid the confounding effect from non-confluent mCherry-expressing cells on cell proliferation rate^95^. A Fisher’s exact test was applied to compare the proportion of S-G2M cells between TF and control cells. A Wilcoxon Rank Sum test was applied to compare the distributions of S or G2M scores between TFs and the control. A kernel density estimate and two sample comparison tests were applied to infer and compare the 2D density distributions of S and G2M scores between TF and control cells using the kde.test function of ks^96^. All statistical tests in these analyses were followed by FDR correction. An adjusted *P* value < 0.05 was considered statistically significant.

#### Analyses for TF pair screening

For the experiments containing TF pair cells, cell barcodes commonly detected by the 10x and TF enrichment library were assigned as being a TF pair- or a single TF-overexpressing cell in a two step process. First, the two TF-IDs accumulating the most reads in the TF-ID-enriched library were aggregated as “main pseudo TF-ID”. Cell barcodes with less than 5 reads of “main pseudo TF-ID” were filtered out. Knee point analysis as described in the *10x scRNA-seq data preprocessing and quality control* section was performed on this aggregated value to filter background cell barcodes. A second knee point analysis was then performed on the number of reads of the main TF-ID (accumulating the most reads for each cell barcode) to distinguish singlets (above knee point threshold) from doublets and TF pair cells (below knee point threshold). By design, the TF-IDs used for the TF pair experiments were different from those in the single TF overexpression experiments. Consequently, cell barcodes below the threshold that did not display expected TF-ID combinations were filtered out as potential doublets. Among the remaining cell barcodes, cells were stratified according to them overexpressing a single TF or TF pair based on whether both the first and second TF-ID were in our list of TF-ID combinations. We performed the following analyses separately for each TF pair, where we subsetted the data each time for cells that were assigned as TF1+TF2, TF1, TF2, or mCherry from the knee point analysis. We then assigned each cell to one of four groups, being TF1+TF2 (>4 UMIs for both TF1 and TF2), TF1 (>4 UMIs for only TF 1), TF2 (>4 UMIs for only TF 2) or control (all other cells). To determine whether the TF pair cells grouped together into a state that was distinct from either the TF1 or TF2 groups, we determined for each cell their 5-nearest neighbors in PCA space (first 20 dimensions). We then quantified for each cell within the TF1+TF2 group the proportion of cells to which it was closest in the other groups, and averaged this over all cells. Cells were binned for both TFs in 4 uniform bins between 0 and the maximal log1p UMI counts with an additional bin for 0 UMI counts.

To detect genes that were uniquely expressed in TF pair cells, we performed differential expression using Seurat’s FindMarkers. Specifically, for each cell within the TF1+TF2 group that had at least 50% TF1+TF2 cells as nearest neighbor, we determined its closest matches to either the TF1 or TF2 groups by performing the 5-nearest neighbor analysis in PCA space (first 20 dimensions), and performing differential expression between the union of these cells with the TF1+TF2 cells. Genes unique to the TF1+TF2 group were defined as those with FDR corrected p-value < 0.05 and absolute fold-change > 1.5.

### Materials availability

Barcoded TF ORF libraries generated by this study are available upon request to W.C. and B.D.

## Data and code availability

- All raw sequencing data generated in this paper has been deposited at ArrayExpress with the accession number <u>E-MTAB-13010</u> and will be publicly available upon publication. Microscopy data reported in this paper will be shared upon request to W.C. and B.D.
- All source code for this paper can be found at https://github.com/DeplanckeLab/TF-seq and is publicly available upon publication.
- Any additional information required to reanalyze the data reported in this paper is available upon request to W.C. and B.D.

## Acknowledgement

We thank Can Aztekin (EPFL), Qiuxia Zhou (EPFL), Olga Pushkarev (EPFL) and Orane Guillaume-Gentil (EPFL) for reviewing the manuscript and providing valuable feedback. We thank Jiahuai Han (Xiamen University) for kindly providing the pBOB-GFP plasmid and Didier Trono (EPFL) for kindly providing the pSIN-TRE-GW-3xHA-puroR plasmid. We thank Horia Hashimi and Marie Rumpler for image analysis support. We thank the Gene Expression (GECF, EPFL) core facility for technical support. This work was supported by Swiss National Science Foundation grants (no. 310030_197082, CRSII5_186271, and TMAG-3_209335) and institutional funding (EPFL) to B.D., a National Key R&D Program of China (grant no. 2021YFA0911100), Shenzhen Medical Research Funding Program (grant no. B2302017) and a Shenzhen Institute of Synthetic Biology Scientific Research Program (grant no. JCHZ20210003) to W.C., a Marie Skłodowska-Curie fellowship (101028476) to W.S., and a Marie Skłodowska-Curie fellowship (101026623) and an EMBO long-term fellowship (2020-895) to G.v.M.

## Author contributions

W.L., W.S., P.R., W.C., and B.D. designed the study. M.B. and W.C. performed experiments with the support of W.L., J.R., and T.L. W.L., W.S., and P.R. performed data analysis with the support of V.G, A.G, and G.v.M. W.L., W.S., P.R., W.C., and B.D. wrote the manuscript with input from M.B., V.G., A.G and G.v.M.

---

**Supplementary Fig. 1:**
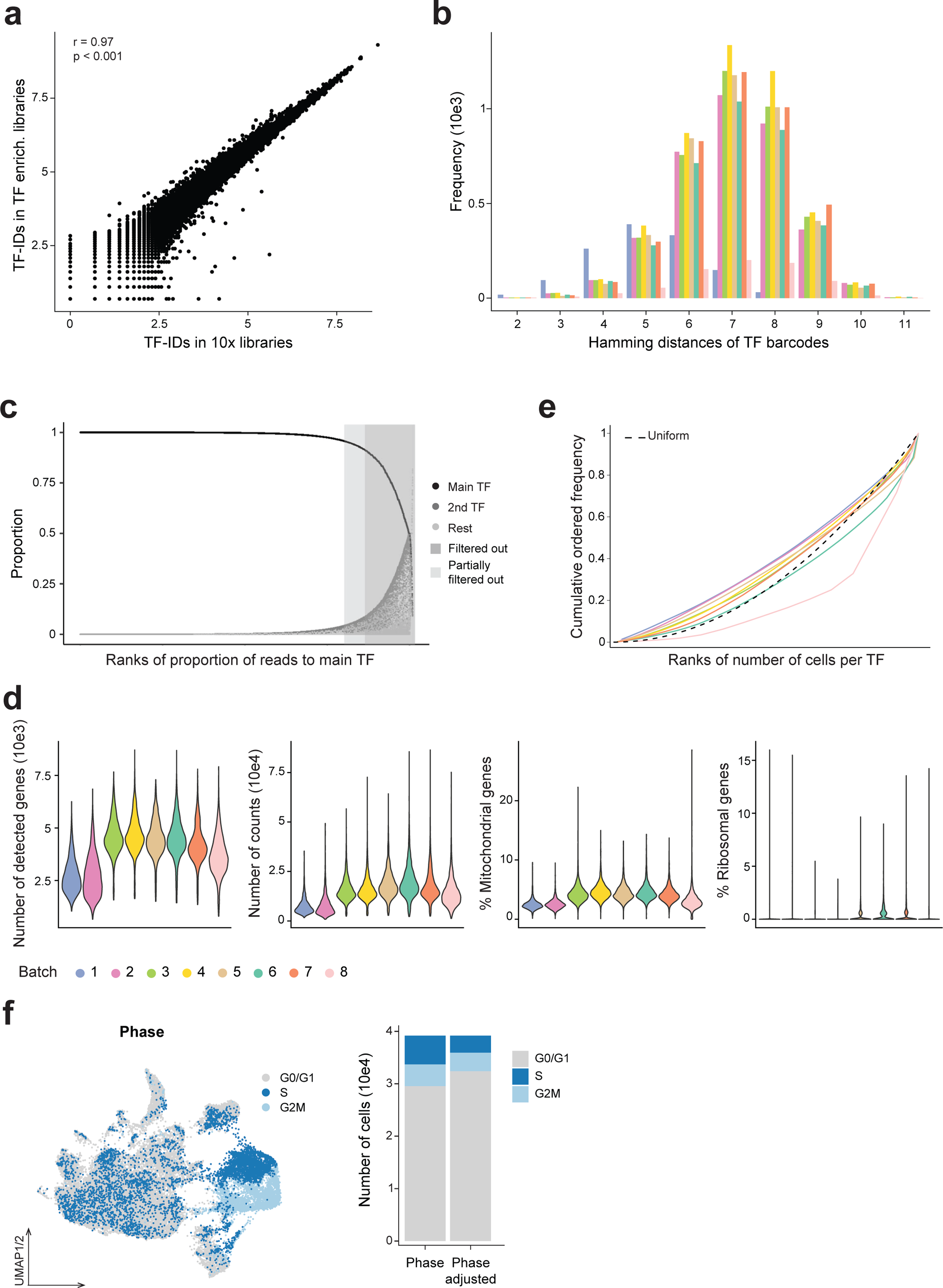
Data quality control and TF assignment, related to Fig. 1. **a** Correlation between the log normalized values of reads mapping to the overexpression construct in the 10x libraries and in the TF enrichment libraries from experiments 1-7 as examples. Pearson correlation coefficient (r) and associated p value are shown. **b** Bar plot showing hamming distances of each pair of TF-ID barcodes and their frequency in each batch indicated by colors. **c** Dot plot showing summarized results of TF assignment for all cells. First, based on the TF-ID count matrix, the proportion of the most detected TF-ID (main TF), the second TF-ID (2nd TF), and the sum of the rest TF-IDs (rest) are calculated for each cell. Cells containing less than 5 counts of TF-IDs are excluded. Then cells in a batch were ordered decreasingly by the proportion of their main TFs, on which a knee point is identified as the threshold to define and filter out cell doublets having a reduced proportion of main TFs in each batch. Finally, cells from all batches are plotted together. The light gray and dark gray zones mark cells partially filtered out (because the threshold is different for each batch) and cells that are all filtered out. **d** Violin plots presenting the number of detected genes and counts, the percentage of mitochondrial and ribosomal gene expression of each cell across batches after quality control. **e** Cumulative ordered frequency of the number of cells assigned to each TF in each batch. The dashed line shows the cumulative ordered frequency of a uniform distribution as comparison. **f** UMAP (left) of the TF atlas colored by cell cycle phase assigned with a default cutoff of 0. Bar plot (right) of the fraction of cells in each phase for both default (0) and adjusted (0.1) cutoffs (see also the UMAP of the TF atlas colored by adjusted cell cycle phase in Fig. 5a and **Methods**).

**Supplementary Fig. 2:**
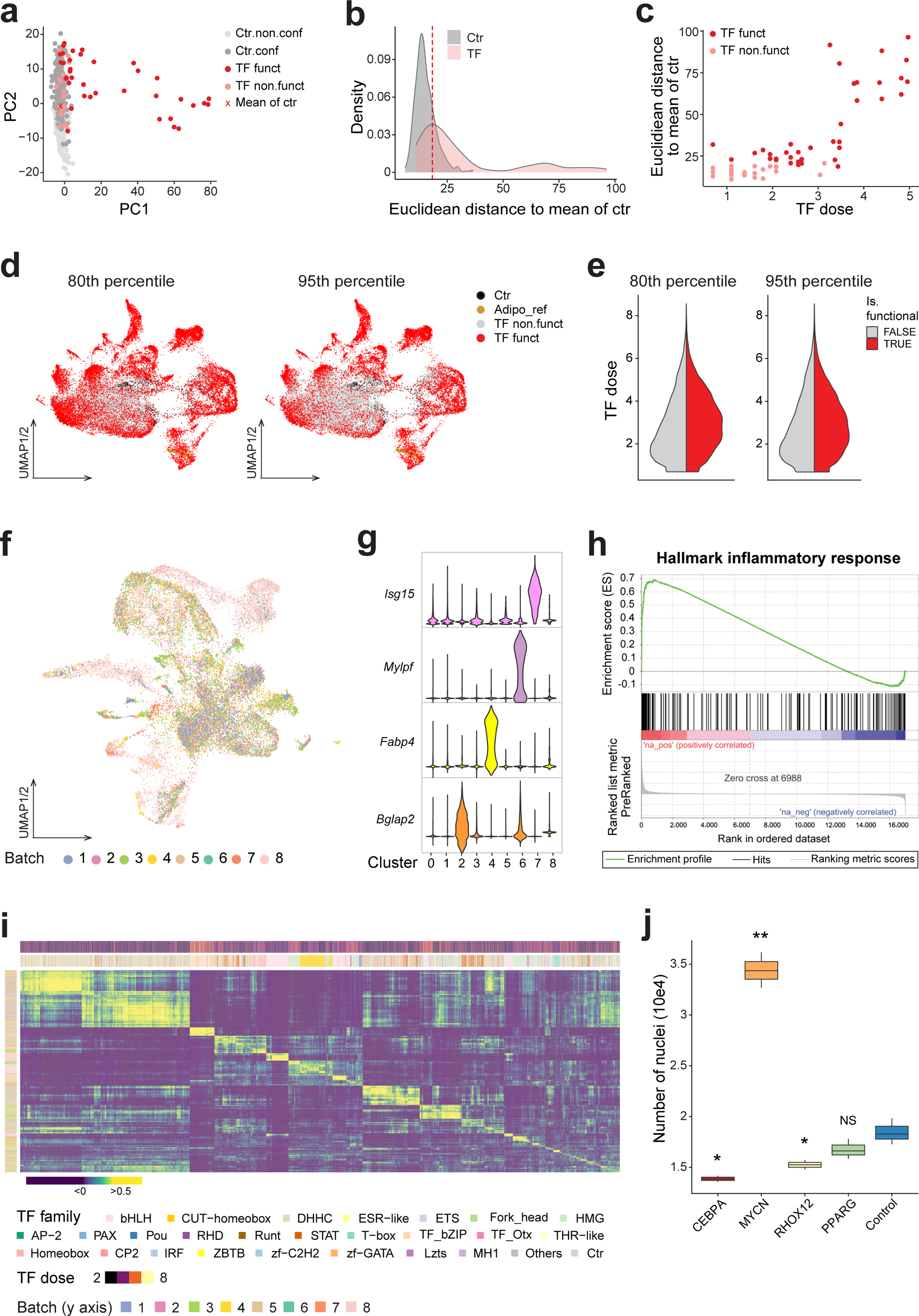
Identification of functional TF cells and lineage-specific clusters, functional similarity of TF families, related to Fig. 2. **a-c** An example of identifying functional TF cells. PCA analysis (**a**) was performed on CEBPA cells of one batch and their batch-paired control (ctr) cells involving both non-confluent (Ctr.non.conf) and confluent (Ctr.conf) mCherry expressing MSCs, allowing to calculate the Euclidean distance between each cell to the mean of control cells of interest in the respective PCA space. CEBPA cells having a distance larger than the distance that 80% of control cells have to their mean (**b**) were selected as ‘functional’ (funct) CEBPA cells. The Euclidean distance of functional and non-functional CEBPA cells was plotted over TF, here *Cebpa*, dose (**c**). **d, e** UMAP of the TF atlas (**d**). Colors indicate control (ctr), adipocyte reference (Adipo_ref), functional (funct) and non-functional TF cells identified using the 80th percentile (left) or 95th percentile (right) as respective thresholds. Violin plots (**e**) showing TF dose in functional (red) or non-functional (gray) cells of all TFs using the two thresholds. **f** UMAP of the functional TF atlas colored by batch. **g** Expression of marker genes for myogenic (*Mylpf*), osteogenic (*Bglap2*), or adipogenic (*Fabp4*) lineages, or immune response (*Isg15*). **h** GSEA result of a hallmark inflammatory response performed between cluster 7 *versus* cluster 0 (containing most of the control cells) in the functional TF atlas (see Fig. 2b). **i** Heatmap showing a pairwise Pearson correlation of cells annotated by TF family, TF dose (in column), and batch (in row). Only TF families that have at least 30 cells were plotted here. Cells are ordered by hierarchical clustering. **j** Nuclei counts quantified on images shown in Fig. 2e. * *P* value < 0.05; ** *P* value < 0.01; NS, not significant; pairwise two-sided *t*-test followed by FDR correction.

**Supplementary Fig. 3:**
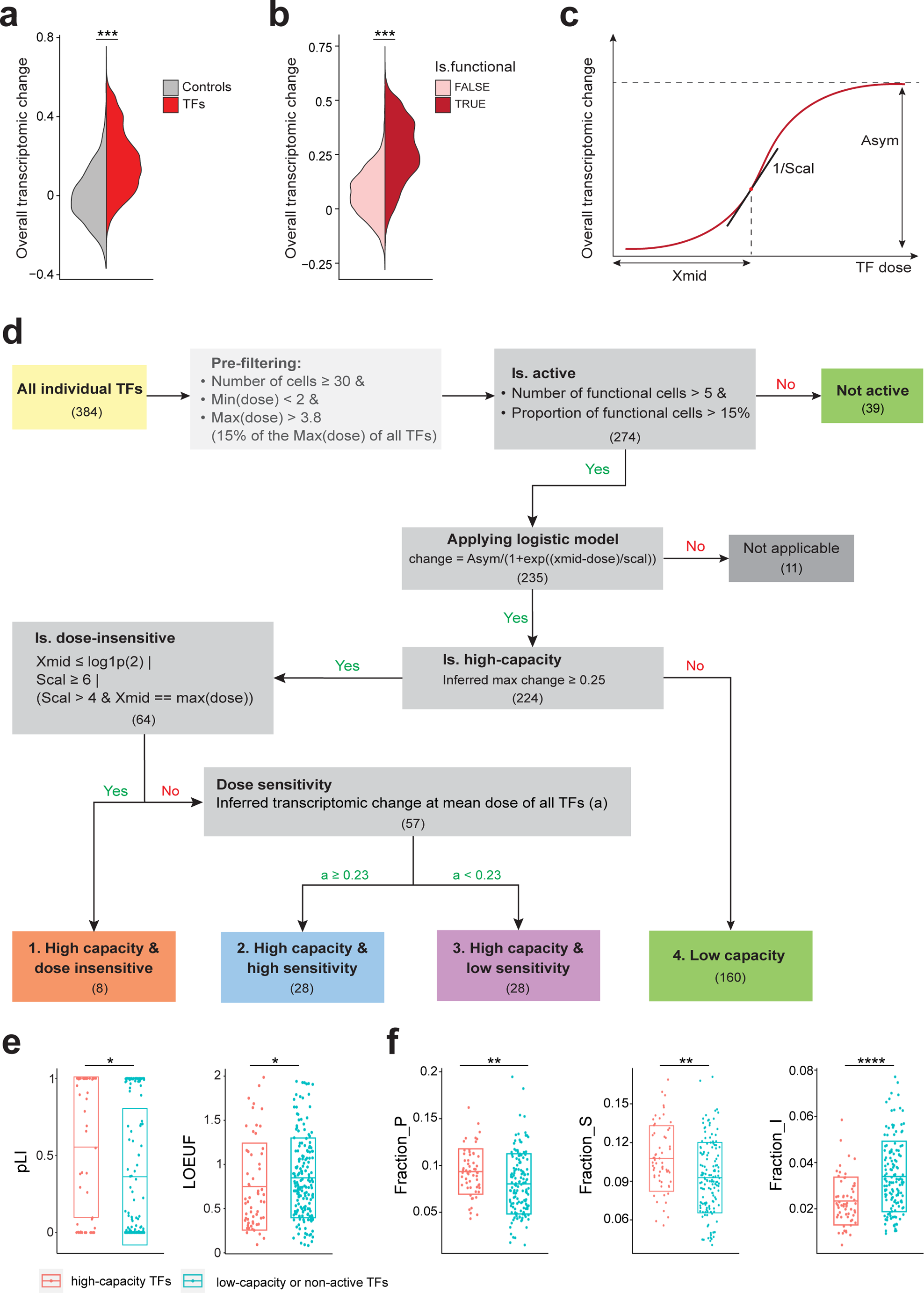
Categorizing TFs based on dose sensitivity and reprogramming capacity, related to Fig. 3. **a, b** Violin plots showing overall transcriptomic change grouped by (**a**) control (mCherry) and TFs cells, or (**b**) functional and non-functional TF cells. **c** Schematic of the logistic regression model on the overall transcriptomic change over TF dose. Asym, Xmid, and Scal are the three parameters estimated by the logistic model used to classify TFs into each category, as described in **d**. **d** Flowchart showing categorization of TFs based on their dose sensitivity and reprogramming capacity by using the logistic regression models (see also **Methods**). **e** Boxplots showing the pLI (probability of being loss-of-function intolerant) scores and the LOEUF (loss-of-function observed / expected upper bound fraction) across human orthologs of high-capacity TFs and low-capacity or non-active TFs. The recommended cutoffs 0.9 and 0.6 were applied to pLI and LOEUF, respectively. pLI > 0.9 or LOEUF < 0.6 represents that a TF is constrained, indicative of a deleterious functional effect. \**P* value ≤ 0.05, Fisher’s exact test. **f** Boxplots showing the distribution of various TF features (Fraction_P, fraction of proline; Fraction_S, fraction of serine; Fraction_I, fraction of isoleucine) across high-capacity TFs and low-capacity or non-active TFs. ** *P* value ≤ 0.01, **** *P* value ≤ 0.0001, Wilcoxon Rank Sum test followed by FDR correction.

**Supplementary Fig. 4.1:**
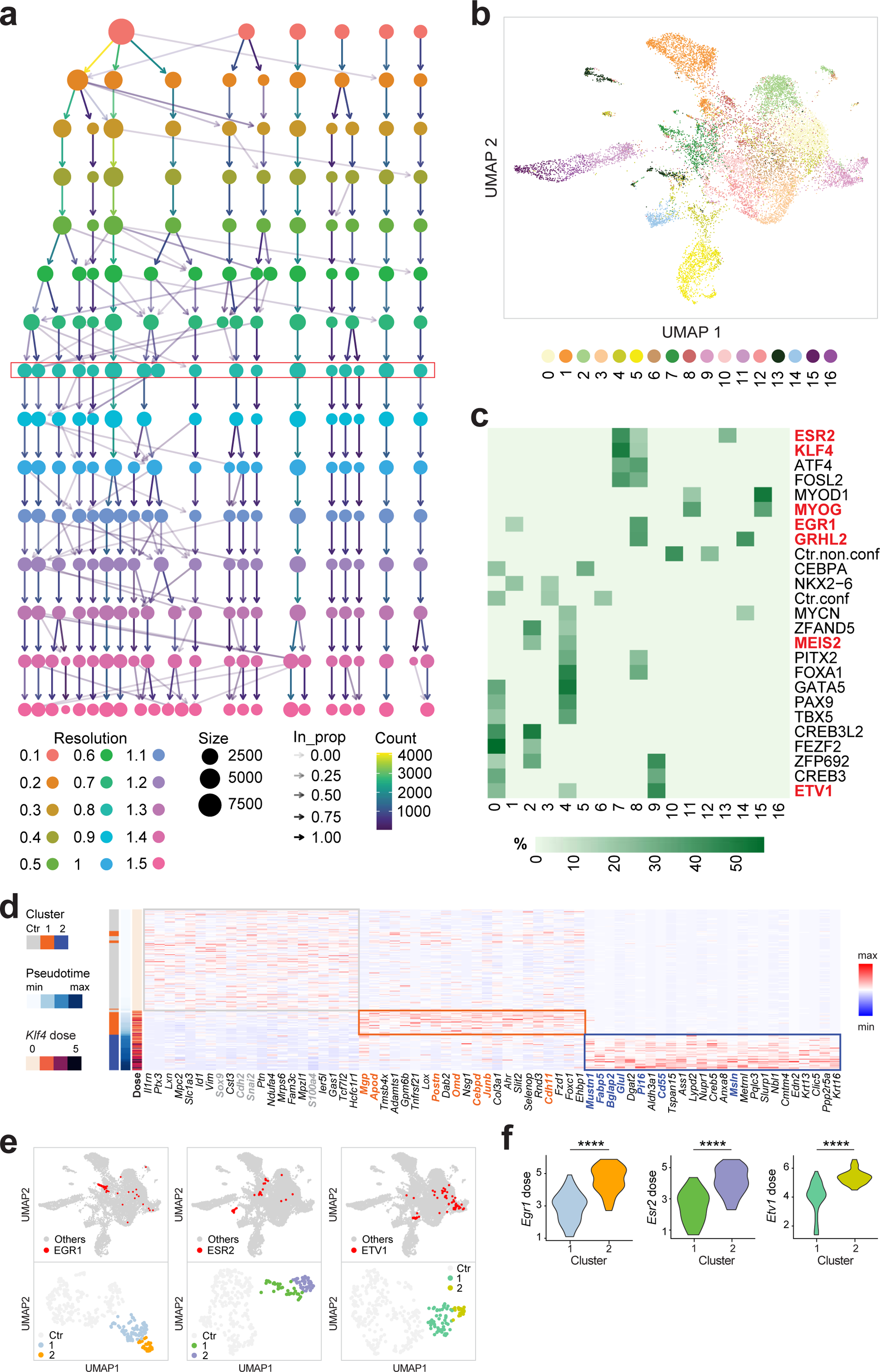
Discovery of candidate TFs associated with cell fate branching, related to Fig. 4. **a, b** Clustering tree (**a**) of the Seurat-based clustering result of the functional TF atlas (as shown in Fig. 2a, b) with only G1 (default phase) cells (**b**), visualizing the relationships between clustering at different resolutions. The resolution 0.8 was used for the functional atlas in **b** to generate an optimal number of clusters such that known lineage cells are separated. **c** Heatmap showing the proportion of TF cells in each cluster relative to the total number of TF cells. Ctr.conf, confluent control cells; Ctr.non.conf, non-confluent control cells. Only controls and TF candidates identified by cell fate branching analysis (see **Methods**) were plotted. **d** Heatmap displaying log-normalized expression (Z-score scaled by gene) of the top markers of GRHL2 clusters (shown in Fig. 4f). Cells are ordered by inferred pseudotime (see also **Supplementary Fig. 4.2h**). Color bars on the left represent GRHL2 clusters, pseudotime, and *Klf4* dose. **e** UMAP (top) of the G1-phase functional TF atlas (shown in **b**) with EGR1, ESR2, and ETV1 cells highlighted in red, and UMAP (bottom) of cells from these individual TFs and the batch-paired control colored by clusters identified by unsupervised clustering. The clusters containing mainly control cells were merged and labeled as the Ctr cluster. Clusters containing mainly TF cells are highlighted in color. **f** Violin plots showing the distribution of *Egr1*, *Esr2*, or *Etv1* dose in cells from clusters shown in the lower panel of **e**.

**Supplementary Fig. 4.2:**
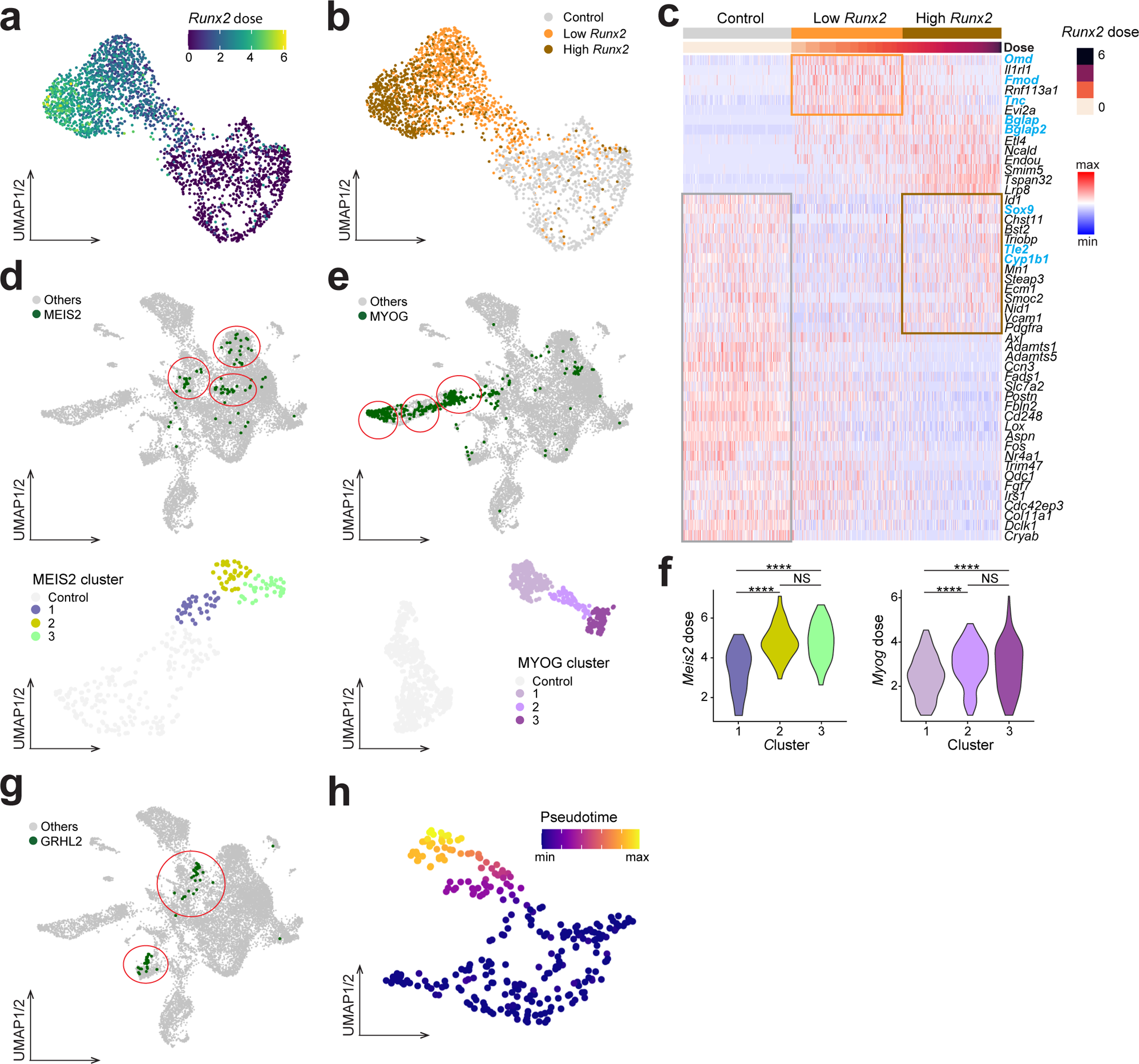
Examples showing dose-dependent or/and stochastic cell fate branching, related to Fig. 4. **a, b** UMAPs of RUNX2 and batch-paired control cells colored by *Runx2* dose (**a**) or groups classified according to *Runx2* dose (**b**). **c** Heatmap displaying log-normalized expression (Z-score scaled by gene) of differentially expressed genes of high and low *Runx2* and control cells. Cells are ordered by *Runx2* dose as indicated by the color bar on the top. **d, e** UMAP (top) of the G1-phase functional TF atlas (shown in **Supplementary Fig. 4.1b**) highlighting cells overexpressing *Meis2* (left) or *Myog* (right). UMAP (bottom) of MEIS2 (left) or MYOG (right), and batch-paired control cells colored by clustering results. The clusters containing mainly control cells were merged and labeled as the Control cluster. Clusters containing mainly MEIS2 or MYOG cells are highlighted in color. **f** Violin plots showing the distribution of *Meis2* (left) or *Myog* (right) dose in the clusters shown in **d** and **e** respectively. **g** UMAP of the G1-phase functional TF atlas highlighting cells overexpressing *Grhl2*. **h** UMAP of GRHL2 and batch-paired control cells colored by inferred pseudotime.

**Supplementary Fig. 5:**
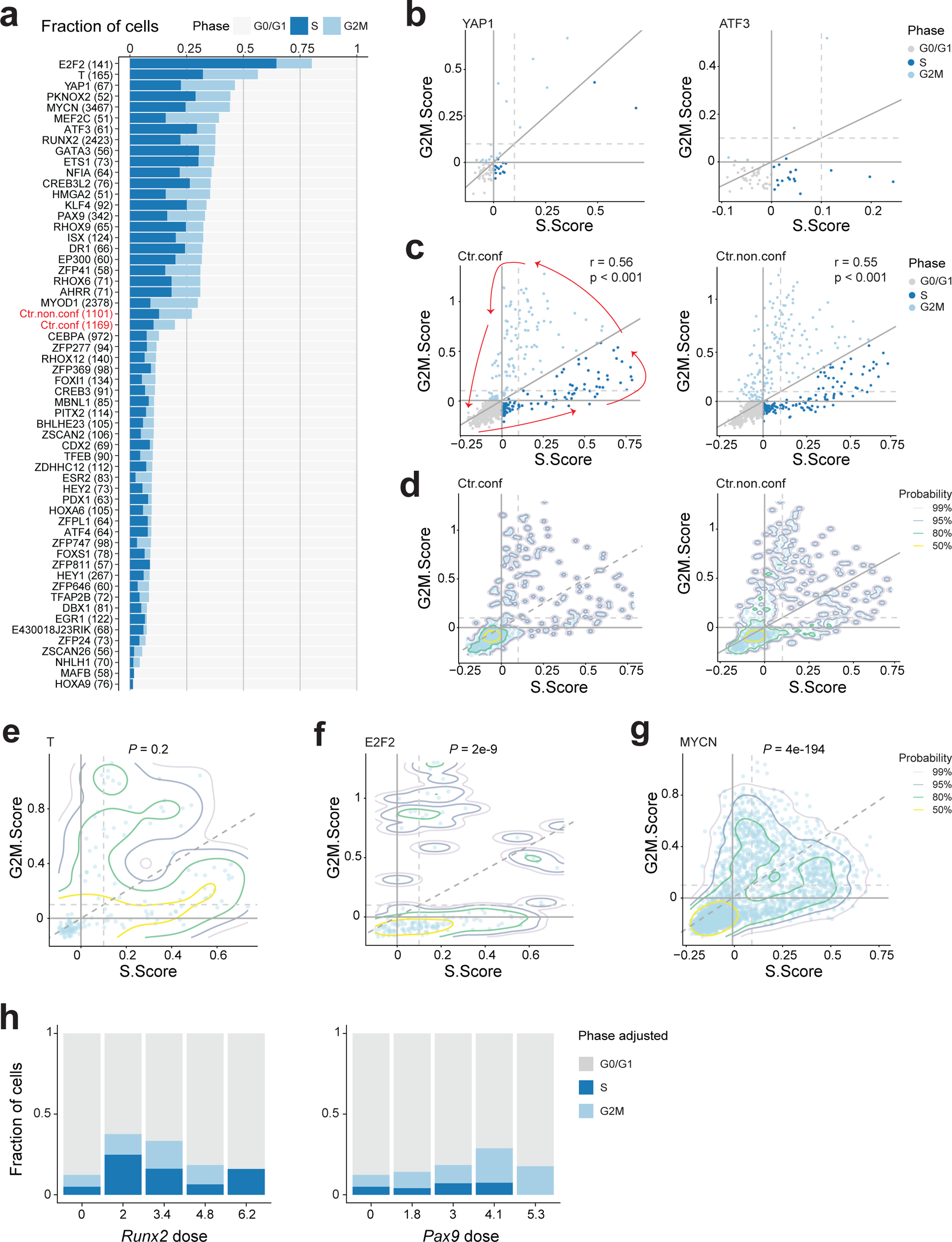
Characterization of cell cycle dynamics regulated by TFs and their doses, related to Fig. 5. **a** Bar plot showing the fraction of cells in the cell cycle phase identified by using the default threshold of cell cycle scores from Seurat. The total number of cells is indicated in brackets. A Fisher’s exact test was performed between confluent control (Ctr.conf) and each TF. Only TFs and non-confluent control (Ctr.non.conf) that tested significantly (FDR-adjusted p-value < 0.05) are visualized here. **b** Scatter plots showing the correlation between S and G2M scores of two TF examples (YAP1 and ATF3), where the fraction of cells in each phase is strongly affected by the selected threshold (0 or 0.1, the latter is indicated by dashed gray lines). Cells are colored by the default phase as in **a**. **c** Scatter plot showing the correlation between S and G2M scores in confluent (left) and non-confluent (right) control cells. Cells are colored by their default cell cycle phase as in **a**. The Pearson correlation coefficient (r) and associated p value are shown. Red arrows on the left indicate the direction of cell cycle progression. **d-g** 2D density contour plots of S and G2M scores showing cell cycle dynamics of confluent (**d**, left) and non-confluent control (**d**, right), T (**e**), E2F2 (**f**), and MYCN (**g**) cells. *P*, adjusted *P* value generated by a two-sample 2D Kernel Density based global comparison test between TF and confluent control followed by FDR correction. The colors of contour lines represent the probability associated with the computed highest density region. For example, 50% represents the region capturing 50% of the data points. **h** Bar plots showing the fraction of cells in each adjusted cell cycle phase across binned doses of *Runx2* or *Pax9*.

**Supplementary Fig. 6:**
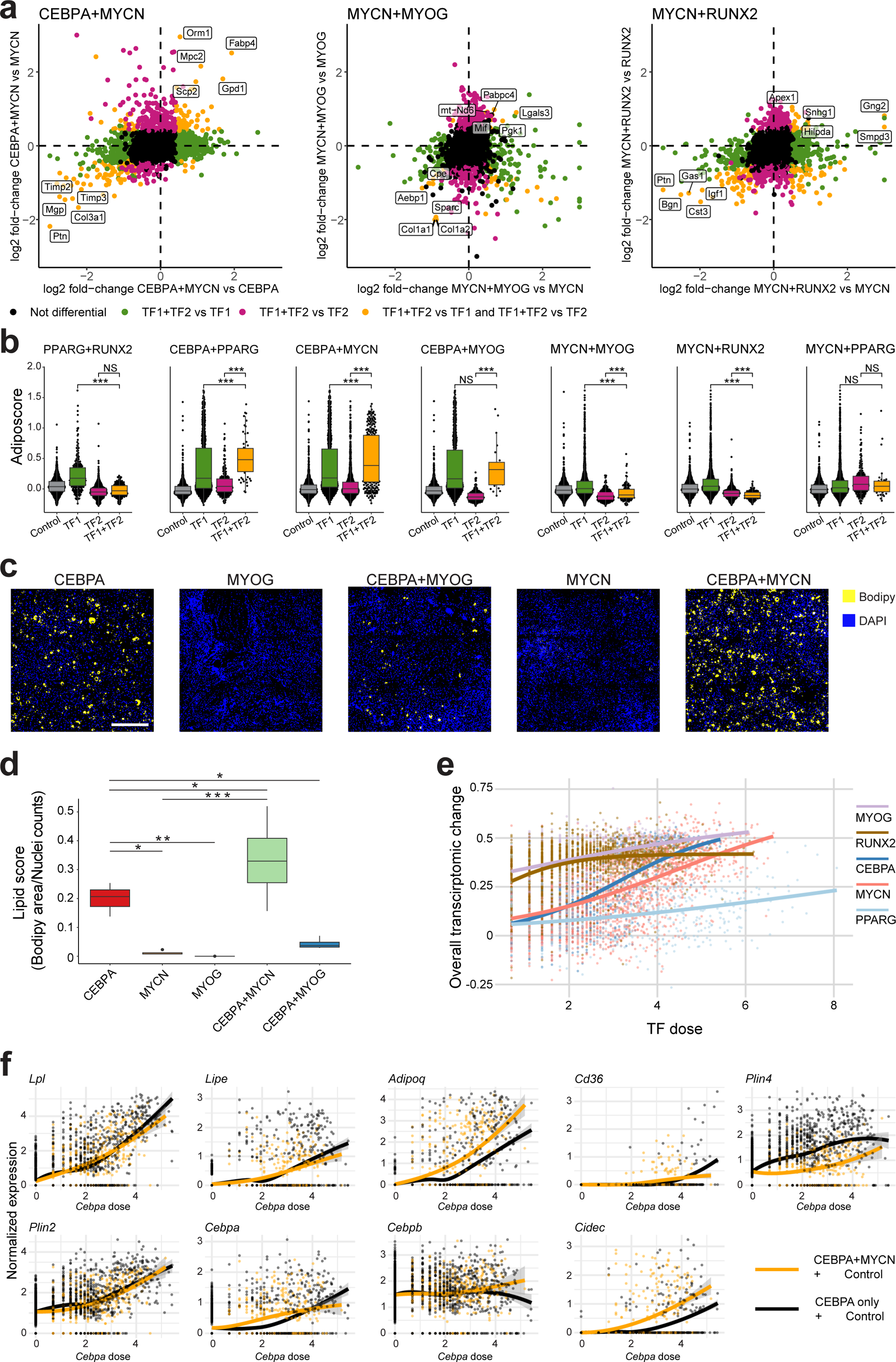
Global and dose-dependent transcriptomic and phenotypic interactions between pairs of TFs, related to Fig. 6. **a** Differential expression between cells with a pair of overexpressed TFs compared to those with a single TF overexpression. Shown are log2-fold changes between TF1+TF2 cells compared to a set of reference cells, defined as the union of all 5-nearest neighbors single TF cells for each TF1+TF2 cell. Significant differential expression was defined using an FDR-adjusted p-value < 0.05 and absolute log2 fold-change > 1.5. Highlighted in orange are those genes that were up- or down-regulated in both comparisons, and that constitute genes that are uniquely regulated in the combination cells. **b** Transcriptomic adiposcore between control cells, single TF cells and TF1+TF2 cells. *** *P* value < 0.001; NS, not significant; Wilcoxon rank sum test followed by FDR correction. **c** Representative fluorescent images of lipid droplets (stained with Bodipy, yellow) and nuclei (stained with DAPI, blue) in single TF and TF1+TF2 cells. Scale bar, 500 μm. **d** Quantification of the lipid score as in (**c**). At least 3 biological replicates for each group. *** *P* value < 0.001; ** *P* value < 0.01; * *P* value < 0.05, pairwise two-sided *t*-test followed by FDR correction. **e** Scatter plot showing the overall transcriptomic change across TF dose colored for the individual TFs involved in combinatorial experiments. The lines represent the fitted logistic regression. **f** Dose response-curves for all transcriptomic adiposcore genes in control and CEBPA cells (black), compared to those for control and MYCN*+*CEBPA cells (orange). The band represents the 95% confidence interval on the smoothened mean using locally estimated scatterplot smoothing.

